# Transport of Alzheimer’s Associated Amyloid-β Catalyzed by P-glycoprotein

**DOI:** 10.1101/2020.10.22.350777

**Authors:** James W. McCormick, Lauren Ammerman, Gang Chen (党陈), Pia D. Vogel, John G. Wise

## Abstract

P-glycoprotein (P-gp) is a critical membrane transporter in the blood brain barrier (BBB) and is implicated in Alzheimer’s disease (AD). However, previous studies on the ability of P-gp to directly transport the Alzheimer’s associated amyloid-β (Aβ) protein have produced contradictory results. Here we use molecular dynamics (MD) simulations, transport substrate accumulation studies in cell culture, and biochemical activity assays to show that P-gp actively transports Aβ. We observed transport of Aβ40 and Aβ42 monomers by P-gp in explicit MD simulations of a putative catalytic cycle. In *in vitro* assays with P-gp overexpressing cells, we observed enhanced accumulation of fluorescently labeled Aβ42 in the presence of Tariquidar, a potent P-gp inhibitor. We also showed that Aβ42 stimulated the ATP hydrolysis activity of isolated P-gp in nanodiscs. Our findings expand the substrate profile of P-gp, and suggest that P-gp may contribute to the onset and progression of AD.

## INTRODUCTION

Alzheimer’s disease (AD) is a progressive and irreversible neurodegenerative disease that primarily affects geriatric populations. One of the pathological hallmarks of AD is deposition and accumulation of amyloid-β (Aβ) in the brain, which is thought to be caused by decreased clearance of Aβ from the brain [1, 2]. Consequently, AD can be diagnosed by the presence of extracellular neurofibrillary tangles composed of Aβ polymers in post-mortem brain sections [3]. While the polymers of Aβ that make up the neurofibrillary tangles are insoluble and stable, the monomeric form of Aβ is soluble [4, 5]. The most common Aβ isoform in the brain is the 40 residue peptide Aβ(1-40); however, in certain forms of AD, the 42 residue isoform Aβ(1-42) has also been shown to increase significantly in the brain. Aβ(1-42) is also regarded as the more fibrillogenic of the peptides produced from amyloid precursor protein (APP) degradation [6–8].

There is compelling preclinical evidence that the blood-brain barrier (BBB), a structure that maintains homeostasis in the central nervous system and protects the brain from harmful substances, plays an important role in Aβ clearance. The BBB is a complex structure composed of endothelial cells of the capillary beds of the brain as well as other supporting cells. Joined by tight junction complexes, these capillary endothelial cells express numerous transport proteins and efflux pumps, thereby creating a selectively permeable barrier that serves to sequester the brain from the rest of the body [9]. To be cleared from the brain, Aβ must therefore cross the BBB. There is evidence that Aβ efflux across the BBB is a multistep process involving several cofactors, in which the LDL Receptor Related Protein 1 (LRP-1) mediates Aβ uptake at the abluminal surface of brain capillary endothelial cells [10–12]. It has been suggested that Aβ is transferred from LRP-1 to P-glycoprotein (P-gp, ABCB1) in endosomes with the help of PICALM and Rab11, and then actively exported by P-gp at the luminal surface from the endothelium into the blood [10–13]. Although experimental results are mixed, there is evidence that P-glycoprotein actively participates in the efflux of both Aβ(1-42) and Aβ(1-40) (Aβ42 and Aβ40, respectively) [10, 14–21].

P-glycoprotein is a member of the ATP Binding Cassette (ABC) transporter family and effluxes a variety of substrates, including drug-like compounds of various sizes and chemical properties [20]. The expression of P-gp in brain capillaries is inversely correlated with deposition of Aβ in the brain, a hallmark of AD pathophysiology [10, 22]. Furthermore, endothelial BBB expression of P-gp declines as humans age, and this decrease in P-gp expression is accompanied by reduced functioning of the BBB [23–25]. Taken together, these data suggest an active role of P-gp in Aβ clearance from the brain. This hypothesis is strengthened by comparing cognitively normal brains to age-matched brains of AD patients; the brains of AD patients exhibit significant decreases in P-gp expression and significant increases in Aβ deposition [23, 26]. Mouse models of AD support an active role for P-gp in the transport of AD-associated Aβ [5, 12, 16, 18, 20, 23, 26].

However, *in vitro* studies of interactions between P-gp and Aβ have yielded mixed results. In 2001, Lam et al. observed that Aβ interacts with and is transported by hamster P-gp [27]. Two subsequent studies, one using human colon adenocarcinoma cells, the other using P-gp transfected porcine LLC cells, support these findings as well [20, 28]. Additionally, inhibition of P-gp in the human hCMEC/D3 cell line resulted in increased intracellular accumulation of Aβ40 [29]. In contrast, a study using paired P-gp expressing and P-gp overexpressing human carcinoma lines found that Aβ42 had no effect on the efflux of a P-gp substrate [14]. This study also found that Aβ42 had no effect on the ATPase activity of P-gp in membrane vesicles [14]. Lastly, the overexpression of P-gp in polarized canine MDCK cells did not promote the transcytosis of radiolabeled Aβ40 in transwell assays [13].

In this study, we assessed the ability of P-gp to transport Aβ using both computational and *in vitro* techniques. Using explicit all atom MD simulations, we analyzed and modeled the transport mechanism of Aβ40 and Aβ42 by human P-gp. In biochemical assays, we showed that Aβ42 stimulates the ATPase activity of purified P-gp; however, this stimulation is dependent upon the lipid environment used. In assays performed in cell culture, which were designed to show potential accumulation of transport substrates (cellular accumulation assays), we observed enhanced retention of fluorescently-labeled Aβ42 in the presence of Tariquidar, a potent P-gp inhibitor [30]. To our knowledge, we are the first to use MD simulations to study the transport of Aβ by P-gp. We are also the first to assess the intracellular accumulation of Aβ42 using a specific P-gp inhibitor in paired P-gp overexpressing and non-overexpressing human cell lines. Our results indicate that Aβ is a transport substrate of P-gp, suggesting that P-gp may be involved in the onset and progression of AD. Understanding the role of P-gp in AD may be of crucial importance for the development of future treatments, and may have implications for compounds targeting P-gp in cancer treatment.

## RESULTS

### Targeted Molecular Dynamics Simulations Show Transport of Amyloid-β by P-glycoprotein

P-glycoprotein (P-gp) is an efflux transporter that is highly promiscuous with respect to transport substrates; P-gp has transmembraneous Drug Binding Domains and two cytoplasmic Nucleotide Binding Domains, at which ATP hydrolysis occurs (Figure 1) [31]. Upon binding of a transport substrate and ATP, the NBDs associate and the DBDs transition from open-to-the-inside (cytoplasm) to open-to-the-outside (extracellular space), resulting in transport of the substrate across the cell membrane and in ATP hydrolysis [31]. Both Aβ-42 (4514.04 Da) and Aβ-40 (3429.80 Da) are significantly larger than the largest known substrate of P-gp, cyclosporine A (1,202.61 Da) [32–34]. To investigate whether P-gp is indeed capable of transporting these bulky Aβ peptides, we performed targeted molecular dynamics (TMD) experiments using techniques that have been previously used to study the transport of P-gp substrates [35, 36].

**Figure 1.**
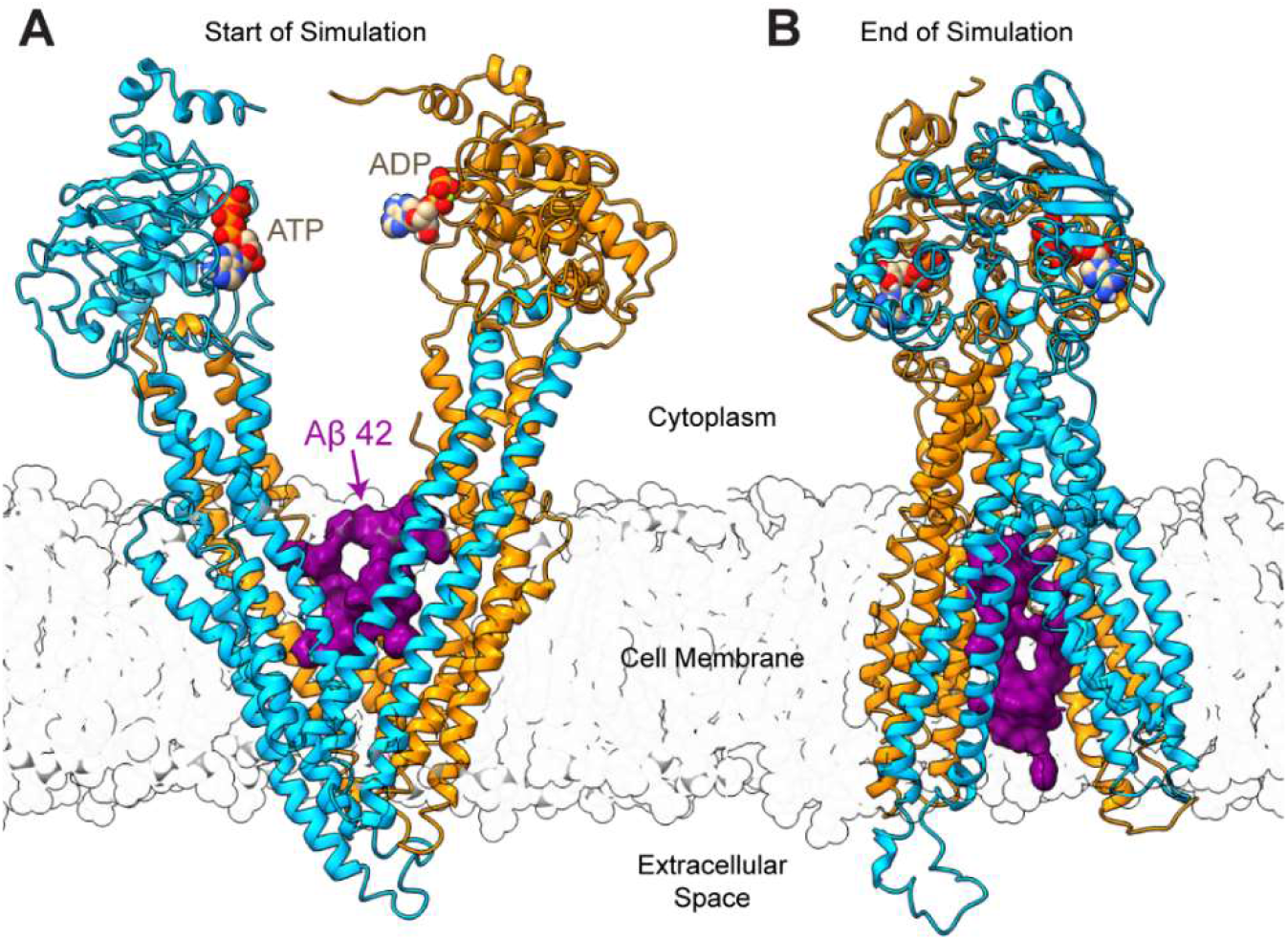
First and Final Frames of Targeted Molecular Dynamics (TMD) Experiments - P-glycoprotein (P-gp) with bound Amyloid-β Peptide: **A)** The first frame of a representative simulation of Aβ42 (PDB 1IYT) bound to the drug binding domains of P-gp. **B)** The final frame of the same representative TMD simulation shown in **(A).** The N- and C-terminal halves of P-gp are colored turquoise or orange; Aβ42 is shown in purple surface representation; ATP and ADP are bound at the nucleotide binding domains and shown in van der Waals representation.

To generate a plausible starting point for TMD simulations, three variants of AD-associated Aβ (Aβ40: PDB IDs 2LFM, 2M4J; Aβ42: PDB ID 1IYT) were docked to human P-gp in a putative starting conformation with the drug binding domains (DBDs) open to the cytoplasm [4, 31, 35, 37–39] (Figure 1, Figure S1), see Methods. When docked to the DBDs, each Aβ peptide occupied the previously observed R and H drug binding sites simultaneously, and Aβ-protein contacts were dominated by hydrophobic interactions (Figure S2 A-C, Table S1) [35, 40]. As a potential negative control, we mutated every residue in the Aβ42 peptide (1IYT) into arginine, creating polyarginine 42 (P-42). P-42 does not fall into the category of compounds normally transported by P-gp, most of which are hydrophobic [41], but it is similar in molecular weight and size to the tested Aβ peptides (Figure S2D). After assembly of a complete system with Aβ, P-gp, lipids, water and ions, each system was relaxed in unbiased molecular dynamics (MD) simulations [35, 36]. TMD simulations were then performed as described in McCormick et al 2015 [35, 36]. Briefly, small forces were applied to α-carbons of P-gp to guide the protein through a series of conformational changes, thereby modeling a putative catalytic transport cycle (Figure S1). Except for the α-carbons of P-gp, no external forces were applied to the Aβ peptides or to any other atoms in the systems.

In each TMD simulation starting with P-gp open to the cytoplasm and with Aβ bound at the DBDs, we observed vectorial movement of Aβ perpendicular to the membrane and towards the extracellular space (n=6 independent simulations per ligand) (Figure 2, Figure S2E-H). The P-gp – bilayer system was oriented such that the membrane is parallel to the X – Y plane, and movement through the membrane is indicated as movement along the Z-axis. In each set of simulations, we observed movement of the Aβ peptide from the cytoplasmic leaflet of the membrane to the extracellular leaflet of the membrane (Figure 2A-C).

**Figure 2.**
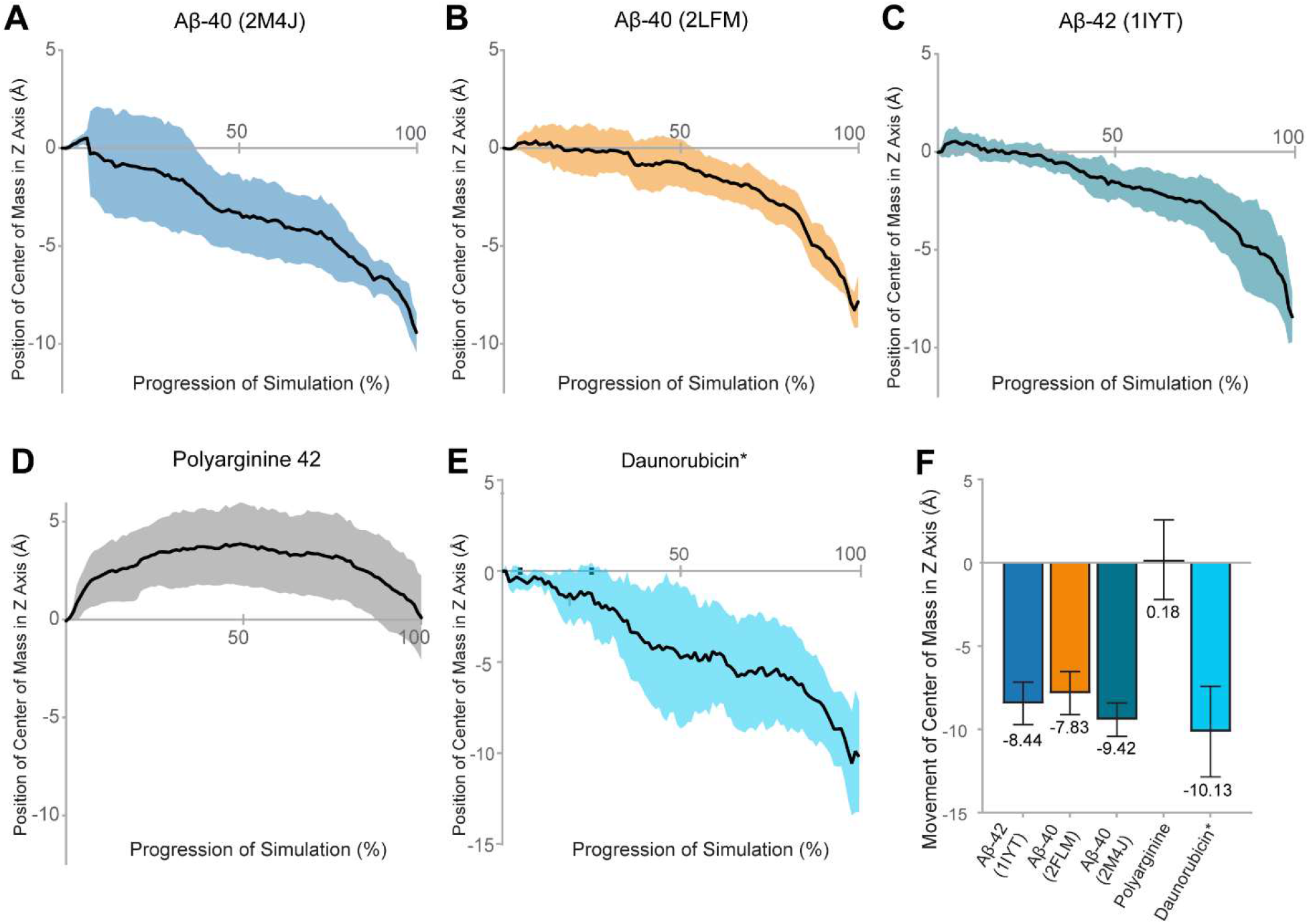
Transport of Amyloid-β by P-glycoprotein in Molecular Dynamics Simulations: The center of mass of **A)** Aβ40 structure 2M4J, **B)** Aβ40 structure 2LFM, **C)** Aβ42 structure 1IYT, **D)** Polyarginine 42 peptide and **E)** daunorubicin were calculated for each step of the simulated putative catalytic cycle of P-gp. Positional changes were calculated relative to the distance from starting coordinates of the ligand. Data represent the mean position of the center of mass (black line) ± one standard deviation from the mean shown in colored shading, n = 6 simulations per ligand. In these simulations, movement towards the cytoplasm is positive on the Z axis, and movement towards the extracellular space is negative. **F)** shows the total mean distance traveled through the plane of the membrane (Z axis) ± one standard deviation. *Data for daunorubicin is reproduced from *McCormick et al. 2015* with permission.

In these simulations, the center of mass of Aβ40 (derived from 2LFM) was observed to move an average of approximately 7.8 ± 1.3 Å from the cytoplasmic to the extracellular side of P-gp (Figure 2A); Aβ40 (from 2M4J) moved an average of approximately 9.4 ± 1.0 Å (Figure 2B); Aβ42 (from 1IYT) moved an average of approximately 8.4 ± 1.3 Å (Figure 2C). For reference, the previously reported transport of daunorubicin (DAU), a known P-gp substrate, is shown in Figure 2E [35]. Daunorubicin was reported to move an average of 10.0 ± 2.7 Å through the DBDs towards the extracellular space [35]. Transport of the Aβ peptides by P-gp ranged from approximately 8 Å to 10 Å across all simulations.

Although both structures of Aβ40 (2LFM and 2M4J) were transported in our simulations, we observed a significant difference between the distance traveled by the two structures of Aβ40 (P = 0.0388). There was no significant difference between the distance traveled by Aβ42 and either form of Aβ40 (Aβ40 2LFM vs. Aβ42, P = 0.4289; Aβ40 2M4J vs. Aβ42, P = 0.17). Despite the large discrepancy in size (DAU 527.5 g/mol, vs. 4514.04 Da Aβ42), there was no significant difference in the distance traveled by daunorubicin (DAU) and the distance traveled by the Aβ monomers (Aβ42 vs. DAU, P = 0.1956; Aβ40 2LFM vs. DAU, P = 0.0894; Aβ40 2M4J vs. DAU, P = 0.5573). These data suggest that the substrate profile of P-gp may include much larger molecules than was previously thought.

### P-gp does not Transport Polyarginine 42 in TMD Simulations

Simulations with Polyarginine 42 (P-42) were started at the initial docking pose of Aβ42 (1IYT) (Figure 1D). Figure 2D shows the average distance traveled by the center of mass of P-42 during the transport cycle. In stark contrast to the behavior of the Aβ peptides, P-42 was not transported through the membrane bilayer (n=6 independent simulations) but remained at a relatively stable position within the DBDs throughout each simulation (0.2 ± 2.6 Å). We observed a highly significant difference between the distance traveled by P-42 and the distance traveled by any of the Aβ monomers, with P < 0.0001 for all three comparisons, respectively (Figure 2F).

### ATPase Activity of P-glycoprotein in the Presence of Amyloid-β

To test whether Aβ42 interacts directly with P-gp, we used a series of *in vitro* ATPase assays of purified murine P-gp, which has 87% sequence identity and high functional similarity to human P-gp [42]. P-gp exhibits a relatively low rate of ATP hydrolysis in the absence of a transport substrate; the introduction of a transport substrate often results in a several fold increase in ATPase activity [43]. Using ATP hydrolysis assays as described in Brewer at el. 2014, we assessed whether monomeric Aβ42 affects the ATPase activity of P-gp [43, 44]. Purified murine P-gp was functionally reconstituted into mixed micelles or lipid bilayer nanodiscs; the latter are considered a more native-like lipid environment [45]. In these studies, 20μg of P-gp in micelles or 15μg of P-gp in nanodiscs were incubated with Aβ42 (molar ratio of 1:18) with or without 150 μM of verapamil (VPL), a substrate of P-gp.

In mixed micelles (Figure 3, blue bars), the ATPase activity of P-gp was stimulated by verapamil (VPL) as expected. Similar to Bello & Salerno 2015, we found that Aβ42 alone did not stimulate the ATPase activity of P-gp in mixed micelles [14]. Furthermore, a combination of VPL and Aβ42 resulted in an increase of ATPase activity above that observed for VPL alone (P < 0.0079). However, with P-gp in nanodiscs (Figure 3, orange bars), we found that Aβ42 significantly stimulated the ATPase activity of P-gp (P < 0.0024). Interestingly, the combination of VPL and Aβ42 did not significantly stimulate ATPase activity relative to VPL alone. Our data suggest that the effect of Aβ42 on the ATPase activity of P-gp is dependent upon the membrane environment [46, 47]. This could explain the contradictory findings of other ATPase activity studies of Aβ and P-gp; these studies used membrane vesicles derived from a variety of cell types [14, 20, 27]. The stimulation of ATPase activity by Aβ42 indicates that it interacts directly with P-gp and supports the hypothesis that Aβ42 is a transport substrate of P-gp.

**Figure 3:**
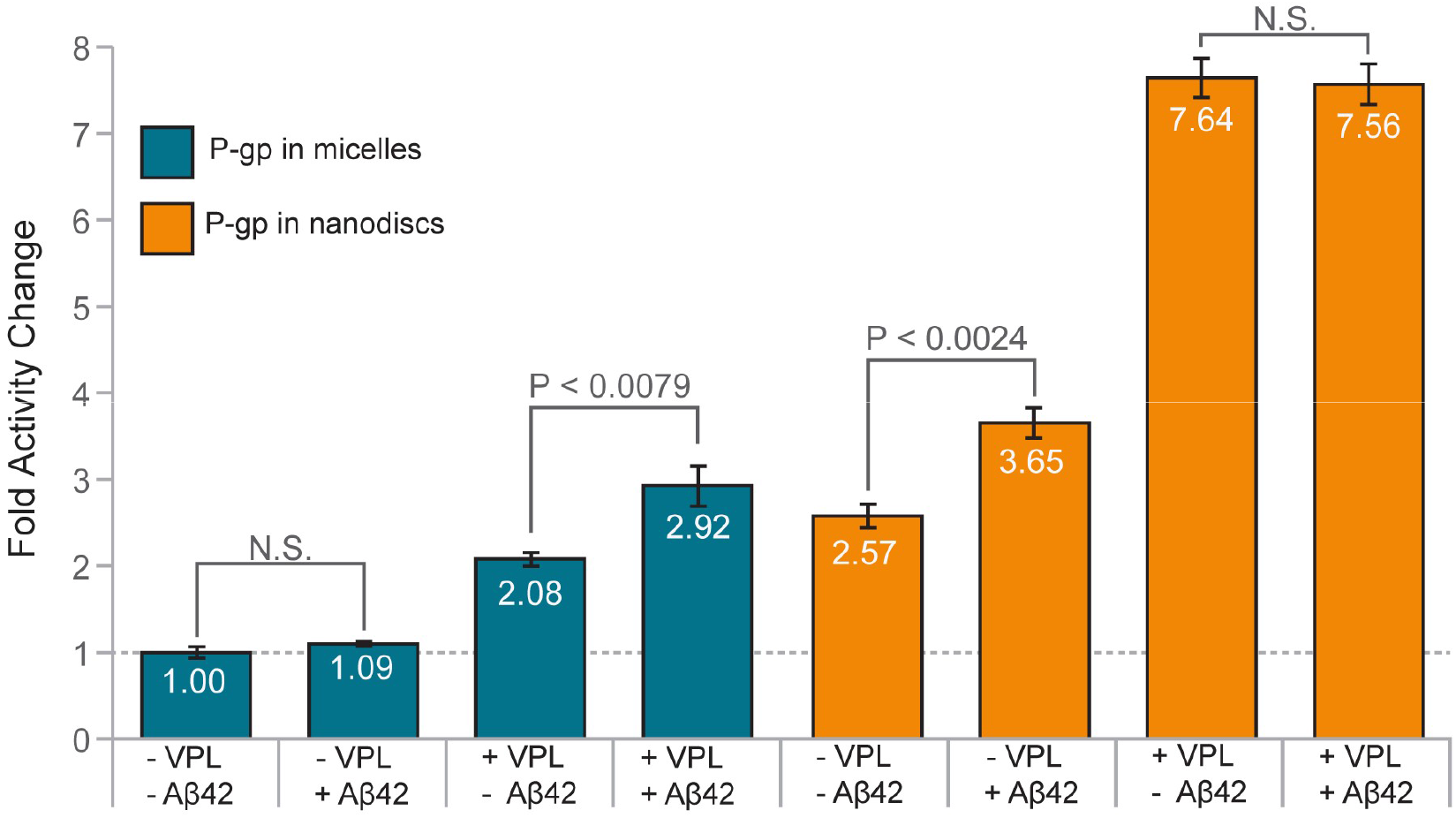
The Effect of Amyloid-β 42 on the ATP Hydrolysis Activity of P-glycoprotein in Micelles and Nanodiscs: The effect of verapamil (VPL) and Aβ42 on the rate of ATP hydrolysis by P-gp was measured in micelles and in nanodiscs. All samples are normalized to the basal ATPase rate of P-gp in micelles (blue bars). Error bars represent ± one standard deviation from the mean.

### Accumulation of Labeled Amyloid-β 42 in DU145-TXR and DU145 Cells

To test whether we could observe the results of transport of Aβ42 by P-gp in human cellular systems, a fluorescently labeled Aβ42 peptide (fl-Aβ42) was assayed for accumulation in the paired DU145 and DU145-TXR prostate cancer cell lines [30, 48, 49]. P-gp is significantly over-expressed in the multidrug resistant DU145-TXR cells relative to the parental, chemotherapy sensitive DU145 cells [49]. In contrast to previous studies of Aβ and P-gp using paired cell lines, we used the strong, selective, and non-competitive P-gp inhibitor Tariquidar (TQR) to assess the accumulation of Aβ [14, 20, 27, 30]. Each cell line was treated with 1μM fl-Aβ42, 1μM of TQR, or a combination of 1μM fl-Aβ42 and 1μM TQR for 16 hours. The accumulation of fl-Aβ42 was quantified using confocal microscopy; Figure 4C-N shows representative images of each treatment (n = 24, two independent trials; see Methods). In both the non-P-gp overexpressing and the P-gp overexpressing cell lines, we observed significant increases (P < 0.0001; P < 0.0001) in the accumulation of fl-Aβ42 in the presence of TQR (Figure 4A-B, Figure S5). Although both cell lines showed a TQR-dependent increase in fl-Aβ42 accumulation, the P-gp overexpressing DU145-TXR cells exhibited the greatest increase in fluorescence (P < 0.0001) (Figure S4). Increased accumulation of Aβ42 upon targeted inhibition of P-gp by TQR indicates that P-gp actively participates in the transport of Aβ42 in human cellular systems.

**Figure 4:**
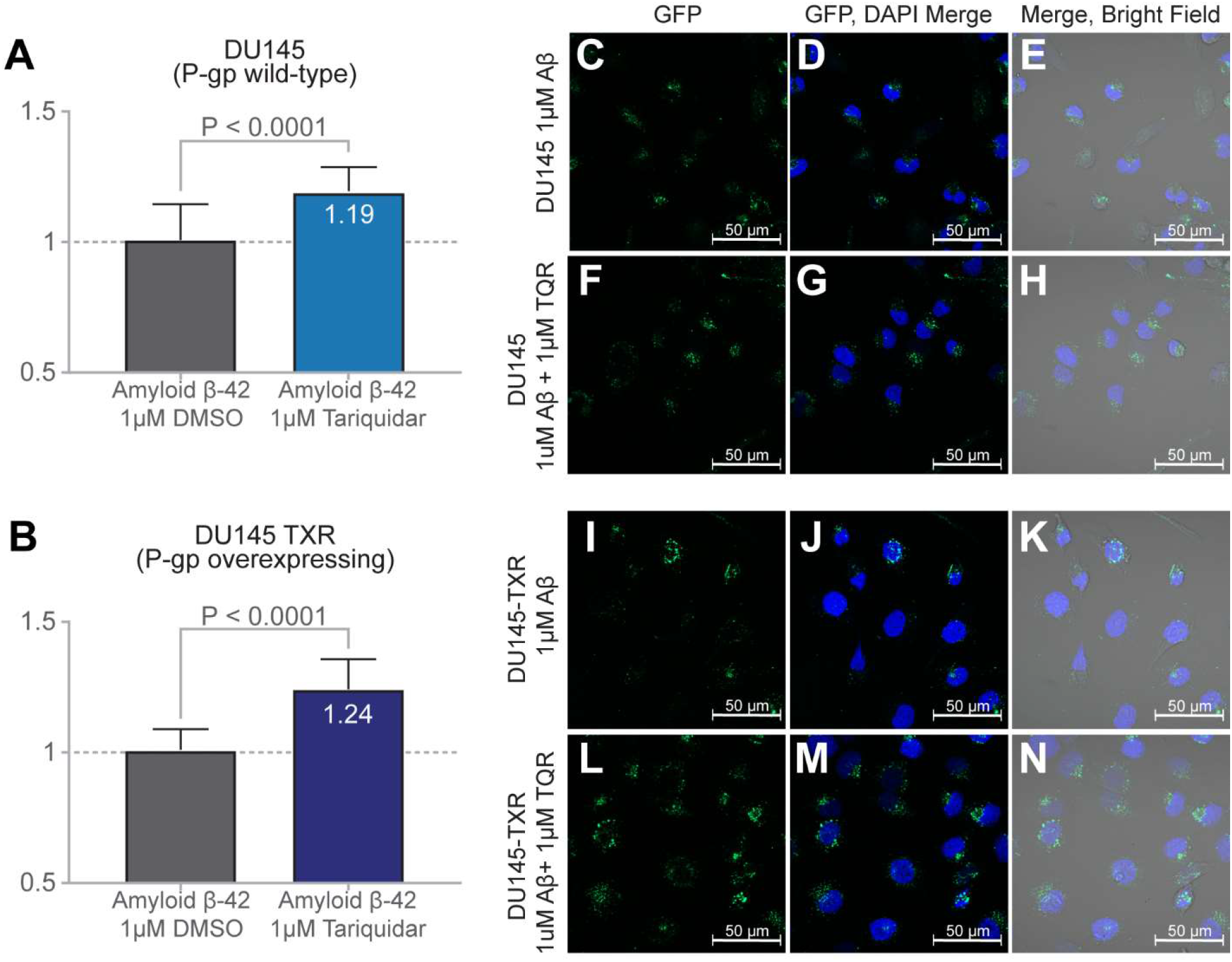
Increased Accumulation of Labeled Amyloid-β 42 in DU145 and DU145TXR Cells after P-gp Inhibition by Tariquidar: The intracellular fluorescence of paired chemotherapeutic sensitive/resistant cancer cell lines (DU145 and DU145-TXR) was measured by confocal microscopy after a 16 hour incubation with fluorescently labeled Aβ42. **A)** When compared to treatment with 1 μM DMSO control (no added TQR, grey bar), wild type P-gp expressing DU145 cells showed a significant 19% (P < 0.0001) increase in mean intracellular fluorescence when inhibited by 1 μM of the P-gp inhibitor Tariquidar (TQR; blue bar, top panel). **B)** Compared to treatment with 1 μM DMSO vehicle (no added TQR, grey bar), treatment of P-gp overexpressing DU145-TXR cells showed a significant 24% (P < 0.0001) increase in mean intracellular fluorescence in the presence of 1 μM Tariquidar (purple bar, bottom panel). Representative Images **C-E** show DU145 treated with 1 μM Aβ42 alone; **F-H** show DU145 treated with 1 μM Aβ42 and 1 μM TQR; **I-K** show DU145-TXR cells treated with 1 μM Aβ42 alone; **L-N** show DU145-TXR treated with 1 μM Aβ42 and 1 μM TQR.

## DISCUSSION

The mechanism by which P-gp might transport substrates of such significant size and flexibility as the Aβ peptides studied here remains unclear. Despite being particularly suited to exploring problems of this nature, MD simulations of Aβ and P-gp have not been performed. Using previously established techniques, we explored how P-gp might transport Aβ40 and Aβ42 using targeted MD simulations. In each simulation of the Aβ peptides, we observed vectorial movement of Aβ through the P-gp DBDs and towards the extracellular space, with total movement ranging between 7.8 and 9.4 Å (Figure 2, Figure S3). These distances correlate well with previously published movements of the P-gp substrate daunorubicin (DAU) [35]. Interestingly, for both Aβ42 and the 2M4J structure of Aβ40, the bulk of observed movement occurred during the transition between the 3B5X and 2HYD conformations of P-gp, or when the DBDs switch from open-to-the-inside to open-to-the-outside (Figure S1, Table S1). In contrast, for both DAU and the 2LFM structure of Aβ40, the bulk of observed movement occurred during the transition from 2HYD to 3B5Z, both of which are open-to-the-outside conformations (Table S1).

Studies have shown that Aβ monomers can fold into structures with two β strands; these β strands allow the monomers to oligomerize and then to potentially assemble into AD-associated amyloid fibrils [50]. At the start of our simulations, the Aβ monomers were not in this folded conformation. It was, however, interesting to ask whether P-gp could somehow facilitate the folding of Aβ monomers during the transport process, and thus contribute to the formation of extracellular amyloid fibrils. Both Aβ40 and Aβ42 stabilize the turn between β strands through a salt bridge between Asp23 and Lys28 [51]. In the simulations reported here, this salt bridge did not form for any significant amount of time (Figure S5). It is possible that hydrophobic interactions with the DBDs prevented the formation of any stable secondary structure by the Aβ peptides (Table S1). Indeed, residue contacts between the Aβ peptides and P-gp were dominated by hydrophobic, non-polar interactions throughout the transport process, with a notable increase in polar contacts as the DBDs opened to the extracellular space (Figure S8, Table S1). Contacts between Aβ and charged residues of the DBDs contributed only a minority of the protein-ligand interactions. Our data suggest that transport by P-gp does not stabilize or contribute to folding of the Aβ monomers. Since Aβ monomers with a distinct folded structure were not simulated, the ability of P-gp to transport or disrupt folded Aβ monomers is unclear and warrants further study.

To date, each study of the effect of Aβ42 upon the ATPase activity of P-gp has used membrane vesicles derived from different cellular systems [14, 27]. Our data suggest that the lipid environment strongly affects the behavior and transport activity of P-gp. Monomeric Aβ42 does not stimulate the ATPase activity of P-gp in micelles but does stimulate the ATPase activity of P-gp in nanodiscs (Figure 3). It has been shown that the structure and behavior of P-gp is affected by the lipid environment [47, 52, 53]. Therefore, we hypothesize that the differences between our tested systems, and potentially the conflicting results of previous studies, may be due to interactions between P-gp the different lipid environments.

Our ATPase studies suggest that Aβ42 interacts directly with P-gp. However, we observed that the Aβ42-stimulated activity was less than half of the ATPase activity stimulated by verapamil (VPL). A possible explanation is that the large size of the Aβ42 peptide (4514 Da for Aβ42 versus 454.6 Da for verapamil) may make difficult for P-gp to move between structural states during the transport process. It is also possible that Aβ42 disassociates slowly from the DBDs due to strong hydrophobic interactions, thus explaining the lower stimulation of ATPase activity compared to VPL (Figure 4, Table S2). While increased stimulation of ATPase activity is considered a characteristic of P-gp substrates, it should be noted that some non-substrates can also stimulate the ATPase activity of P-gp [46, 47, 54].

To test if P-gp can transport Aβ42, we performed fluorescence accumulation assays in a human cellular system. Our data show that inhibition of P-gp by the strong and P-gp specific inhibitor, tariquidar (TQR), resulted in increased intracellular accumulation of fluorescently labeled Aβ42 (fl-Aβ42). This increased accumulation was observed in both P-gp overexpressing DU145-TXR cells and the parental, non-P-gp overexpressing DU145 cells (Figure 4). P-gp is greatly overexpressed in DU145-TXR cells relative to the parental DU145 cells [49]. Interestingly, the uninhibited DU145-TXR cells exhibited significantly higher fluorescence than the uninhibited DU145 cells (Figures 4 and S5). We hypothesize that this is due to reduced CD33 levels in the DU145-TXR cell line (3.75 fold decrease relative to DU145 cells), a byproduct of generating resistance through exposure to the chemotherapeutic Paclitaxel [49]. CD33 is a transmembrane protein involved in cellular adhesion and Aβ clearance processes; reduced expression of CD33 has been shown to result in increased uptake of Aβ42 [55–57]. However, inhibition of the resistant DU145-TXR cells resulted in a 33% greater change in Aβ fluorescence relative to the change in DU145 cells, again strongly suggesting that inhibition of P-gp transport resulted in low levels of Aβ efflux through P-gp with a concomitant increase in Aβ accumulation in these cells.

## CONCLUSION

Through the combination of computational simulations, kinetic measurements of the purified protein, and transport assays in a human cellular environment, we have shown here that P-gp is able to transport the monomeric form of Aβ. While there is a growing body of evidence that P-gp plays an important role in the clearance of Aβ across the BBB, such conclusions are beyond the scope of this study [16, 18, 58]. Adding Aβ to the list of known substrates indicates that P-gp can transport much larger molecules than was previously thought. Given the clinical importance of P-gp and of other ABC transporters, we believe that the ability of human efflux pumps to transport large ligands, and the mechanism by which they do so, warrants further study.

## EXPERIMENTAL PROCEDURES

### Docking Amyloid-β to Human P-glycoprotein (P-gp)

Aβ structures were docked to the drug binding domains (DBDs) of P-glycoprotein in the open-to-the-cytoplasm conformation (derived from the homologous 4KSB structure) of human P-gp using AutoDock Vina as described previously (see Figure S1) [31, 36, 37, 59]. Ligand interactions were limited to the cytoplasmic extensions of the transmembrane helices and the transmembrane sections of P-gp and used an exhaustiveness of 128 (the default exhaustiveness or number of replica docks for Vina is set at 8). Ligand binding to nucleotide binding domains (NBDs) was not investigated. The resultant ligand docking positions were ranked by predicted binding affinities; the conformational pose with the highest predicted affinity was used as a starting point for molecular dynamics (MD) simulations, except where indicated in the text.

Three different structures of Aβ were docked to the DBDs of P-gp. 2LFM, a partially folded solid state NMR structure of Aβ40 in an aqueous environment, docked with a predicted affinity of −7.2 kcal/mol (Figure S2A) [39]. 2M4J, a 40 residue Aβ fibril derived from AD brain tissue docked with a predicted affinity of −7.1 kcal/mol (Figure S2B) [4]. 1IYT, a solid state NMR structure of Aβ42 in an apolar microenvironment, with a predicted affinity of −7.2 kcal/mol (Figure S2C) [4]. As a control, every residue in the highest affinity docking pose of Aβ40 fibril 2LFM was mutated into an arginine, creating the Polyarginine 42 peptide (Figure S2D).

### Transport of Amyloid-β through P-gp in Molecular Dynamics Simulations

To facilitate the efflux of substrates across the cell membrane, ATP-driven ABC-transporter proteins undergo large conformational changes powered by the binding and hydrolysis of ATP. These conformational changes switch the drug binding domains (DBDs) of the transporter from “open to the cytoplasm” (inward facing) to “open to the extracellular space” (outward facing) [60, 61]. Such cycling has been hypothesized for P-gp and has previously been shown by us in computational simulations to transport small-molecule, drug-like ligands from the cytoplasmic membrane leaflet to the extracellular leaflet and extracellular space [35]. The modeled, putative catalytic cycle of P-gp therefore reflects the hypothesized sequence of conformational changes for ABC transporters and has allowed us to visualize substrate transport driven by P-glycoprotein. These previous studies have allowed us to investigate P-glycoprotein-driven movement of small drug-like molecules across the membrane. These computational simulations have been extended here to the larger, polypeptide substrates, Aβ40 and Aβ42.

To model a putative catalytic transport cycle of P-gp, we used crystal structures of P-gp homologues in various conformations as in McCormick et al. 2015. Because these structures were determined from crystals, the conformations represented by these structures are relatively stable, representing relatively low energy conformations of the protein. During TMD simulations, small forces were applied to selected Cα atoms of P-gp to direct the movement of protein domains toward the respective target coordinates. The putative catalytic transport cycle of P-gp we used follows the sequence of conformational states based on earlier work [35, 36]: (1) a conformation with the DBDs wide open to the cytoplasm (derived from 4KSB); (2) a conformation with the DBDs slightly open to the cytoplasm (derived from 3B5X); (3) a conformation with fully engaged NBDs and DBD opened to the exterior (derived from 2HYD); and (4) a final conformation with NBDs in an ATP hydrolysis transition state and the DBDs fully open to the extracellular space (derived from 3B5Z) (see Figure S1). Using these TMD simulations, we guided P-gp through a putative transport cycle and included Aβ40 (2LFM, 2M4J) or Aβ42 (1IYT) in the DBDs on the cytoplasmic side of the membrane [4, 36, 38, 39].

The inward facing structure of the mouse P-gp (4KSB) has fully disengaged NBDs with its transmembrane DBDs oriented in an inward facing state [31]. The structure of MsbA from *Vibrio cholerae* (3B5X) has disengaged NBDs, and DBDs partially open to the cytoplasm. The structure of SAV1866 from *S. aureus* (2HYD) has engaged NBDs and its DBD open to the outside [60, 62]. The structure of MsbA from *S. typhimurium* (3B5Z) also has fully engaged NBDs with its DBD open to the outside but may represent a post-hydrolysis transition state since crystallization conditions included an MsbA - ADP-vanadate complex. Using the structures in the aforementioned sequence, we previously simulated the conformational changes of a putative catalytic transport cycle using models of human P-gp and targeted molecular dynamics (TMD) simulations [35, 36] based on structures from [63, 64].

### Preparation of the Amyloid-β 42 Synthetic Peptide

*Monomerization of Aβ42 for ATPase Assays.* The Aβ42 synthetic peptide was purchased from GenicBio Limited, PRC (sequence DAEFRHDSGYEVHHQKLVFFAEDVGSNKGAIIGLMVGGVV-IA). The peptide had a molecular weight of 4514.14 g/mol and was judged to be 95.40% pure by HPLC. To monomerize the protein, lyophilized Aβ42 was removed from storage at −80°C and allowed to equilibrate at room temperature for 30 minutes to avoid condensation upon opening the vial. In a fume hood, 1 mg of the lyophilized Aβ42 peptide was resuspended in 300 μl of 1,1,1,3,3,3-hexafluoro-2-propanol (HFIP, Sigma-Aldrich). The mixture was sonicated and vortexed thoroughly to ensure proper solvation. The solution was then centrifuged at 10,000 rpm for 5 min and the supernatant was moved to a clean tube. Centrifugation was repeated once more to remove any insoluble materials. The solution was then aliquoted into separate vials where the HFIP was allowed to evaporate in the fume hood overnight. The desiccated pellets were stored at −20°C. For use in ATPase assays, the samples were resuspended in a 1:4 mixture of dimethyl sulfoxide (DMSO) and sterile water.

#### Preparation of the Amyloid-β 42 for Accumulation Assays in Cell Culture

Fluorescent (HiLyte™ Fluor 488) labeled Aβ42 was purchased for cell culture assays from Anaspec (sequence: HiLyte™ Fluor 488 – DAEFRHDSGYEVHHQKLVFFAEDVGSNKGAIIGLMVGGVVIA). This peptide had a molecular weight of 4870.5 g/mol, absorption/emission wavelengths of 503/528 nm, and was judged to be >= 95% pure by HPLC (CAT# AS-60479, LOT# 1958003). To prepare the Aβ peptide for cell culture assays, 0.1mg of the peptide was dissolved in 30 uL of 1% (w/v) ammonium hydroxide in sterile water and filtered using a 0.45 uM pore filter [65]. Once thoroughly dissolved and mixed, 380 uL of Phosphate Buffered Saline (PBS) solution were added to the peptide-ammonium hydroxide solution (final NH4OH 0.07%). After mixing thoroughly again, the peptide solution was aliquoted into 50 μL aliquots and frozen until use. When using the peptide solution for cell culture experiments, the peptide containing aliquot was thawed and vortexed immediately before use.

### Accumulation of Labeled Amyloid-β 42 in DU145 and DU145-TXR Cells

Multidrug-resistant (MDR) DU145-TXR prostate cancer cells have been previously shown to overexpress P-glycoprotein [49]. These MDR DU145-TXR cells (kindly provided by Evan Keller, Univ. of Michigan) were derived from drug sensitive DU145 cancer cells by culturing in the presence of the chemotherapeutic paclitaxel to create the P-gp overexpressing cell line DU145-TXR [49]. Both DU145 and DU145-TXR cells were grown in complete media consisting of RPMI-1640 with L-glutamine, 10% fetal bovine serum, 100 U/mL penicillin and 100 μg/mL streptomycin in a humidified incubator at 37 °C using 5% CO2. The drug-resistant line DU145-TXR was maintained under positive selection pressure by supplementing complete media with 10 nmol/L paclitaxel. Both cell lines were grown and seeded on collagen-treated flasks and plates (Collagen Type I, Corning).

To assess the accumulation of fluorescently labeled Aβ in both cell lines, cells were trypsinized from monolayers and seeded at 60,000 cells per well in 6 well plates in complete RPMI media. Prior to seeding, a sterilized glass coverslip was placed in each well, and each well was treated with a working solution of 0.01 mg/mL collagen Type I (Corning) in 0.02 N Acetic Acid for 10 minutes and rinsed twice with PBS. After 24 hours incubation at 37 °C, the media was removed and replaced with fresh complete RPMI media. The cells were dosed with 1 μM of the P-gp inhibitor tariquidar (TQR), 1uM of Hylyte 488 Aβ42, a combination of both, or DMSO as a control, and incubated at 37 °C and 0.05% CO_2_ for 16 hours [66]. The final concentration of ammonium hydroxide in each well was kept at approximately 0.0035 % and was matched in controls to ensure identical treatment of all cell samples. If there was any leftover Aβ in the thawed aliquot, the excess Aβ was not re-frozen but was discarded to avoid aggregation that can be induced by freeze-thawing. After 16 hours of incubation, media was removed, and cells were gently washed with cold PBS. Cells were then fixed in 4% paraformaldehyde in PBS for 20 minutes. Cells were then stained with DAPI in PBS for 10 minutes, and subsequently washed twice with cold PBS. Each coverslip was removed from the well and mounted on a glass slide using Fluoromount G mounting fluid.

After drying slides overnight, fluorescence-confocal microscopy was performed on a Zeiss LSM800 microscope using Plan-Apochromat 20x and 40x/1.3 oil-immersion objectives. Images were captured using the Zeiss ZEN software; images taken at 40X were used for quantification of fluorescence. Experiments were performed in duplicate, with two slides per treatment, and in two trials. At 20X magnification, 12 images per slide, two trials (n = 48 per treatment) were taken. At 40X magnification, six images were taken per slide with fl-Aβ42 treatment (n=24 total) for quantification; at least three images were taken for TQR-only or DMSO control slides to confirm the lack of fl-Aβ42 fluorescence seen at 20X magnification. All images were taken before analysis was performed; all images captured at 40X magnification of slides treated with fl-Aβ42 are included in the analysis (see Figure S6 and S7).

Quantification was performed using FIJI ((Fiji Is Just) ImageJ, NIH, Bethesda, Maryland, USA)[67–69]. To quantify fluorescence with ImageJ, raw CZI image files were imported using the Bio-Formats plugin [70]. Analysis of Aβ fluorescence with ImageJ was automated using ImageJ macros (included in Supplementary Files). Using a copy of the raw green channel image, regions of Aβ fluorescence were “thresholded” as follows: (1) use the default “Subtract Background” function (rolling ball radius of 10px); (2) apply the default Unsharp Mask (radius 1 px, mask=0.60 sigma) to define the edges of fluorescent areas; (3) threshold the image (minimum 15, maximum 255) to determine which areas will be measured; (4) convert the thresholded image to a binary mask; (5) save the binary mask as a .TIF image. Through this process, a binary mask is created in which areas of thresholded Aβ fluorescence are black, and all other areas are white. The binary mask was then used to define regions for measurement on the original, raw green channel image. Using the binary mask to define regions for measurement, areas of thresholded Aβ fluorescence in the raw green channel image were measured using the default “Analyze Particles” function of ImageJ (see Supplementary Files). Using the previously created binary mask of each image, the mean background intensity of each raw green channel image was measured by selecting the inverse of the Aβ areas, and then using the default “Measure” function. Aβ Fluorescence is quantified as a measure of the “corrected mean fluorescence intensity” using the following formulae and is reported in arbitrary units.

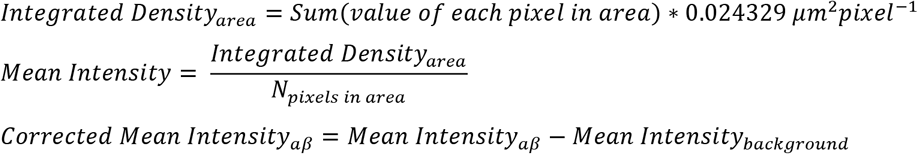

Once the initial analysis was completed using ImageJ, images were examined for potential extracellular aβ fluorescence using the Bright Field channel overlaid with the green channel as a guide. In the event of extracellular particles, we (1) created a copy of the original binary mask for that image; (2) manually removed the extracellular particle from the binary mask copy; (3) ran the measuring and analysis functions using the ‘corrected’ binary mask to redirect measurements. Any extracellular areas were manually removed from the binary mask, and the image was re-analyzed using the same macro. This resulted in negligible changes from the original values and did not affect the overall results of the analysis (data in supplemental). To compare the levels of Aβ fluorescence between treatments, we report the Corrected Mean Intensity in arbitrary units. Data was analyzed using a two-tailed T test with equal variance and GraphPad Prism version 7 for Windows, GraphPad Software, San Diego, California USA, www.graphpad.com.

### ATPase Activity Assays with P-gp in Nanodiscs and Micelles

#### Purification of Murine P-gp

Murine cys-less P-gp was used for all ATPase activity assays [42, 43, 71, 72]. Protein purification of P-gp and ATP hydrolysis assays were performed as described in Delannoy et al 2005 and Brewer et al. 2014 with some modifications as described below [43, 71].

#### Preparation of P-gp in Micelles

P-gp was expressed in *Pichia Pastoris* GS-115. To isolate P-gp from its native membrane and embed it in detergent micelles, 80 mL of frozen cell pellets were thawed in a 37 °C water bath, and protease inhibitors (160 μL pepstatin A, 32 μL Leucine, 16 μL chymostatin, 800 μL of 200 mM PMSF and 800 μL of 200 mM DTT) were subsequently added. Cells were then broken open using 175 mL of glass beads and a BeadBeater (Biospec products). The BeadBeater was filled to the top with buffer containing 30% glycerol, 50 mM Tris, 125 mM NaCl, 10 mM imidazole (pH 8.0). To prevent the samples from overheating during bead beating, ice with rock salt was used. Samples were spun at 10,000 rpm for 30 min at 4°C to remove debris, nuclei, mitochondria and unbroken cells using a Beckman Avanti JXN-26 centrifuge. The supernatants were then subjected to a fast spin at 45,000 rpm for 45 min at 4°C using Beckman Optima XPN-80. The resultant pellets (which contain P-gp) were washed with microsome wash buffer (20% glycerol, 50 mM Tris, pH 7.4), and resuspended in Tris buffer (30% glycerol, 50 mM Tris, 125 mM NaCl, 10 mM imidazole, pH 8.0). Nickel-NTA columns were used to capture P-gp engineered with His-tag.

Microsomes containing P-gp were diluted with Tris buffer (20% glycerol, 50 mM Tris, 50 mM NaCl, 10 mM imidazole, pH 8.0) to 2 mg/mL, and 0.6% n-Dodecyl β-D-maltoside (DDM, w/v) (Sigma-Aldrich) and 0.01% lysophosphatidylcholine (lyso-PC, w/v) were added. Then, the sample solution was sonicated in an ice-cold water bath (model) for 5 cycles of 5 min. Samples were then spun down at 20,000 rpm for 30 min at 4°C using a Beckman Optima XPN-80 centrifuge to remove undissolved microsomal proteins. The supernatants were then applied to a Ni-NTA gravity column (QIAGEN), and incubated for 30 min at 4°C. Flow-through was collected at 2 mL/min, and the column was washed with 20 bed volume of wash buffer (20% glycerol, 50 mM Tris, 50 mM NaCl, 20 mM imidazole, pH 7.5) at 1mL/min, followed by 10 bed volume of the second wash buffer (20% glycerol, 50 mM Tris, 50 mM NaCl, 40 mM imidazole, pH 7.5) at 1 mL/min. Protein was eluted with buffer (20% glycerol, 50 mM Tris, 50 mM NaCl, 300 mM imidazole, pH 7.5) for 3 bed volumes at 0.5 mL/min. Both buffers for the wash and elution were supplemented with 0.6% DDM (w/v) and 0.01% lyso-PC (w/v) if not otherwise specified. The eluates were concentrated to about 150 μL using Amicon 100K centrifugation filters (MilliporeSigma) at 4°C, and stored at −80°C.

#### Reconstitution of P-gp into Nanodiscs

P-gp was incubated with a 10x molar excess of membrane scaffold protein (MSP) and a 500x molar excess of 40% L-alpha-phosphatidylcholine (PC) from soybean (Sigma) for 1 hour at room temperature with gentle agitation to facilitate formation of nanodiscs [73]. Biobeads SM-2 (BioRad) were presoaked in methanol, washed with a large amount of water, equilibrated with equilibration buffer (20% glycerol, 50mM Tris-HCl pH 7.5, 50mM NaCl), and finally added at a ratio of 1.4g/mL to the assembly mix to remove detergent. After addition of Biobeads SM-2, the mixture was incubated for 1.5 hours at room temperature with shaking to remove detergent from the crude nanodisc sample. The Biobeads were removed from the crude nanodiscs by piercing the bottom of the centrifuge tube with a 25 gauge needle and centrifuging at 1000xg for 1 minute. Empty discs were removed by Ni-NTA column chromatography, utilizing the histidine–tag at P-gp. Six bed volumes of column wash buffer (20% glycerol (v/v) 50mM Tris-HCl pH 7.5 at RT, 50mM NaCl, 20mM imidazole) were then applied. Purified nanodiscs were eluted from the column by applying 1 bed volume elution buffer (20% (v/v) glycerol, 50mM Tris-HCl pH 7.5, 4ºC, 50mM NaCl, 300mM imidazole). Samples were analyzed by gradient SDS-PAGE and coupled enzyme assays.

#### ATPase Activity Assays

Briefly, ATP hydrolysis by P-gp was coupled to the oxidation of NADH to NAD^+^ by two enzymes, pyruvate kinase and lactate dehydrogenase, as described in [43]. The coupled enzyme assay cocktail included 50 mM Tris, pH 7.5, 24 mM MgSO_4_, 20mM KCl, 1.94 mM phosphoenolpyruvate) (PEP), 0.058 mg/mL pyruvate kinase, 0.0288 mg/mL lactate dehydrogenase, 1.13 mM NADH, and 4 mM ATP. The absorbance decrease of NADH at 340 nm was recorded using a BioTek Eon plate reader BioTek. The ATPase activity of P-gp was directly correlated with the rate of NADH oxidation. An extinction coefficient of 6220 M^-1^cm^-1^ at 340 nm was used for calculations of NADH oxidation with a measured path length of 0.6 cm. Aβ42 was incubated with P-gp for 30 min at 37ºC before the addition of coupled enzyme assay cocktail. In the activity assays reported here, 15 μg of purified nanodiscs were used, and 20 ug of purified micelles were used.

This work is supported by NIH NIGMS [R15GM09477102] to JGW, SMU University Research Council, SMU Engaged Learning program, the SMU Center for Drug Discovery, Design and Delivery, the SMU Center for Scientific Computing, the Communities Foundation of Texas, and a private gift from Ms. Suzy Ruff of Dallas, Texas.

The authors would like to acknowledge and thank Kelsey Paulhus for her valuable and constructive suggestions and guidance during the planning and development of the microscopy in this work. We would also like to thank Amila K. Nanayakkara for his valuable contributions he made on a pilot study for the accumulation assays in this work. We would like to thank Dr. Kimberly Reynolds (UT Southwestern Medical Center) for reviewing the manuscript and for her advice. We would also like to thank Professor Heng Du (University of Texas, Dallas) for advice and methods for handling Amyloid β preparations and Professor Evan Keller (U. of Michigan) for the DU145 and DU145-TXR cell lines.

The authors declare no competing financial interest.

## Author Contributions

James McCormick

- Original project design and conception, coordination
- Original and updated literature review
- Data quantification and methodology design of computational experiments
- Writing of original manuscript
- Docking of AB isoforms to PGP
- Fluorescently labeling original AB42 used for pilot cell culture assays
- Monomerization of AB42 for ATPase assays
- Making figures
- Running, designing, analyzing simulations
- Manuscript revision

Lauren Ammerman

- troubleshooting, design, and adaptation of accumulation assay with both cell lines, labeled Aβ, and confocal microscopy
- Data quantification and methodology design using FIJI and ImageJ of cell culture data
- Revision, writing, of original manuscript
- Running molecular dynamics simulations
- Analysis of microscopy data, simulation data
- Making figures
- Updating literature review

Mike Chen

- ATPase assay experiments, protein purification, data analysis of ATPase assays
- Quantification of monomerized AB42 for ATPase assays

Pia D. Vogel

- Interpretation of data
- Editing and revisions of the manuscript

John G. Wise

- refinement and design of the study
- experimental manipulation of Amyloidβ peptides
- interpretation of data
- editing and revisions of the manuscript
- final approval of the version to be published

## References

1. LaFerla, F.M., K.N. Green, and S. Oddo, Intracellular amyloid-beta in Alzheimer’s disease. Nat Rev Neurosci, 2007. 8(7): p. 499–509.

2. Murphy, M.P. and H. LeVine, 3rd, Alzheimer’s disease and the amyloid-beta peptide. J Alzheimers Dis, 2010. 19(1): p. 311–23.

3. Snyder, H.M., et al., Alzheimer’s disease research in the context of the national plan to address Alzheimer’s disease. Mol Aspects Med, 2015. 43-44: p. 16–24.

4. Crescenzi, O., et al., Solution structure of the Alzheimer amyloid beta-peptide (1-42) in an apolar microenvironment. Similarity with a virus fusion domain. Eur J Biochem, 2002. 269(22): p. 5642–8.

5. Wang, W., A.M. Bodles-Brakhop, and S.W. Barger, A Role for P-Glycoprotein in Clearance of Alzheimer Amyloid beta-Peptide from the Brain. Curr Alzheimer Res, 2016. 13(6): p. 615–20.

6. Naslund, J., et al., Relative abundance of Alzheimer A beta amyloid peptide variants in Alzheimer disease and normal aging. Proc Natl Acad Sci U S A, 1994. 91(18): p. 8378–82.

7. Mori, H., et al., Mass spectrometry of purified amyloid beta protein in Alzheimer’s disease. J Biol Chem, 1992. 267(24): p. 17082–6.

8. Schmidt, M., et al., Comparison of Alzheimer Abeta(1-40) and Abeta(1-42) amyloid fibrils reveals similar protofilament structures. Proc Natl Acad Sci U S A, 2009. 106(47): p. 19813–8.

9. Daneman, R. and A. Prat, The blood-brain barrier. Cold Spring Harb Perspect Biol, 2015. 7(1): p. a020412.

10. Chai, A.B., et al., P-glycoprotein: a role in the export of amyloid-beta in Alzheimer’s disease? FEBS J, 2020. 287(4): p. 612–625.

11. Ito, S., S. Ohtsuki, and T. Terasaki, Functional characterization of the brain-to-blood efflux clearance of human amyloid-beta peptide (1-40) across the rat blood-brain barrier. Neurosci Res, 2006. 56(3): p. 246–52.

12. Storck, S.E., et al., The concerted amyloid-beta clearance of LRP1 and ABCB1/P-gp across the blood-brain barrier is linked by PICALM. Brain Behav Immun, 2018. 73: p. 21–33.

13. Nazer, B., S. Hong, and D.J. Selkoe, LRP promotes endocytosis and degradation, but not transcytosis, of the amyloid-beta peptide in a blood-brain barrier in vitro model. Neurobiol Dis, 2008. 30(1): p. 94–102.

14. Bello, I. and M. Salerno, Evidence against a role of P-glycoprotein in the clearance of the Alzheimer’s disease Abeta1-42 peptides. Cell Stress Chaperones, 2015. 20(3): p. 421–30.

15. Bruckmann, S., et al., Lack of P-glycoprotein Results in Impairment of Removal of Beta-Amyloid and Increased Intraparenchymal Cerebral Amyloid Angiopathy after Active Immunization in a Transgenic Mouse Model of Alzheimer’s Disease. Curr Alzheimer Res, 2016.

16. Cirrito, J.R., et al., P-glycoprotein deficiency at the blood-brain barrier increases amyloid-beta deposition in an Alzheimer disease mouse model. J Clin Invest, 2005. 115(11): p. 3285–90.

17. Elali, A. and S. Rivest, The role of ABCB1 and ABCA1 in beta-amyloid clearance at the neurovascular unit in Alzheimer’s disease. Front Physiol, 2013. 4: p. 45.

18. Hartz, A.M., D.S. Miller, and B. Bauer, Restoring blood-brain barrier P-glycoprotein reduces brain amyloid-beta in a mouse model of Alzheimer’s disease. Mol Pharmacol, 2010. 77(5): p. 715–23.

19. Jain, S., et al., Pregnane X Receptor and P-glycoprotein: a connexion for Alzheimer’s disease management. Mol Divers, 2014. 18(4): p. 895–909.

20. Kuhnke, D., et al., MDR1-P-Glycoprotein (ABCB1) Mediates Transport of Alzheimer’s amyloid-beta peptides--implications for the mechanisms of Abeta clearance at the blood-brain barrier. Brain Pathol, 2007. 17(4): p. 347–53.

21. van Assema, D.M. and B.N. van Berckel, Blood-Brain Barrier ABC-transporter P-glycoprotein in Alzheimer’s Disease: Still a Suspect? Curr Pharm Des, 2016. 22(38): p. 5808–5816.

22. Vogelgesang, S., et al., Deposition of Alzheimer’s beta-amyloid is inversely correlated with P-glycoprotein expression in the brains of elderly non-demented humans. Pharmacogenetics, 2002. 12(7): p. 535–41.

23. Chiu, C., et al., P-glycoprotein expression and amyloid accumulation in human aging and Alzheimer’s disease: preliminary observations. Neurobiol Aging, 2015. 36(9): p. 2475–82.

24. van Assema, D.M., et al., Blood-brain barrier P-glycoprotein function in Alzheimer’s disease. Brain, 2012. 135(Pt 1): p. 181–9.

25. Deo, A.K., et al., Activity of P-Glycoprotein, a beta-Amyloid Transporter at the Blood-Brain Barrier, Is Compromised in Patients with Mild Alzheimer Disease. J Nucl Med, 2014. 55(7): p. 1106–11.

26. Wijesuriya, H.C., et al., ABC efflux transporters in brain vasculature of Alzheimer’s subjects. Brain Res, 2010. 1358: p. 228–38.

27. Lam, F.C., et al., beta-Amyloid efflux mediated by p-glycoprotein. J Neurochem, 2001. 76(4): p. 1121–8.

28. Abuznait, A.H., et al., Up-regulation of P-glycoprotein reduces intracellular accumulation of beta amyloid: investigation of P-glycoprotein as a novel therapeutic target for Alzheimer’s disease. J Pharm Pharmacol, 2011. 63(8): p. 1111–8.

29. Tai, L.M., et al., P-glycoprotein and breast cancer resistance protein restrict apical-to-basolateral permeability of human brain endothelium to amyloid-beta. J Cereb Blood Flow Metab, 2009. 29(6): p. 1079–83.

30. Fox, E. and S.E. Bates, Tariquidar (XR9576): a P-glycoprotein drug efflux pump inhibitor. Expert Rev Anticancer Ther, 2007. 7(4): p. 447–59.

31. Ward, A.B., et al., Structures of P-glycoprotein reveal its conformational flexibility and an epitope on the nucleotide-binding domain. Proc Natl Acad Sci U S A, 2013. 110(33): p. 13386–91.

32. Amyloid β Protein Fragment 1-42. [cited 2020 14 August 2020]; Available from: https://www.sigmaaldrich.com/catalog/product/sigma/a9810?lang=en&region=US.

33. Amyloid β Protein Fragment 1-40. [cited 2020 14 August 2020]; Available from: https://www.sigmaaldrich.com/catalog/product/sigma/a1075?lang=en&region=US.

34. Information, N.C.f.B., PubChem Compound Summary for CID 5284373, Cyclosporin A. 2020.

35. McCormick, J.W., P.D. Vogel, and J.G. Wise, Multiple Drug Transport Pathways through Human P-Glycoprotein. Biochemistry, 2015. 54(28): p. 4374–90.

36. Wise, J.G., Catalytic transitions in the human MDR1 P-glycoprotein drug binding sites. Biochemistry, 2012. 51(25): p. 5125–41.

37. Trott, O. and A.J. Olson, AutoDock Vina: improving the speed and accuracy of docking with a new scoring function, efficient optimization, and multithreading. J Comput Chem, 2010. 31(2): p. 455–61.

38. Lu, J.X., et al., Molecular structure of β-amyloid fibrils in Alzheimer’s disease brain tissue. Cell, 2013. 154(6): p. 1257–68.

39. Vivekanandan, S., et al., A partially folded structure of amyloid-beta(1-40) in an aqueous environment. Biochem Biophys Res Commun, 2011. 411(2): p. 312–6.

40. Shapiro, A.B. and V. Ling, Extraction of Hoechst 33342 from the cytoplasmic leaflet of the plasma membrane by P-glycoprotein. Eur J Biochem, 1997. 250(1): p. 122–9.

41. Schinkel, A.H. and J.W. Jonker, Mammalian drug efflux transporters of the ATP binding cassette (ABC) family: an overview. Adv Drug Deliv Rev, 2003. 55(1): p. 3–29.

42. Aller, S.G., et al., Structure of P-glycoprotein reveals a molecular basis for poly-specific drug binding. Science, 2009. 323(5922): p. 1718–22.

43. Brewer, F.K., et al., In silico screening for inhibitors of p-glycoprotein that target the nucleotide binding domains. Mol Pharmacol, 2014. 86(6): p. 716–26.

44. Vogel, P.D. and R.L. Cross, Adenine nucleotide-binding sites on mitochondrial F1-ATPase. Evidence for an adenylate kinase-like orientation of catalytic and noncatalytic sites. J Biol Chem, 1991. 266(10): p. 6101–5.

45. McLean, M.A., M.C. Gregory, and S.G. Sligar, Nanodiscs: A Controlled Bilayer Surface for the Study of Membrane Proteins. Annu Rev Biophys, 2018. 47: p. 107–124.

46. Sharom, F.J., ABC multidrug transporters: structure, function and role in chemoresistance. Pharmacogenomics, 2008. 9(1): p. 105–27.

47. Sharom, F.J., Complex Interplay between the P-Glycoprotein Multidrug Efflux Pump and the Membrane: Its Role in Modulating Protein Function. Front Oncol, 2014. 4: p. 41.

48. Stone, K.R., et al., Isolation of a human prostate carcinoma cell line (DU 145). Int J Cancer, 1978. 21(3): p. 274–81.

49. Takeda, M., et al., The establishment of two paclitaxel-resistant prostate cancer cell lines and the mechanisms of paclitaxel resistance with two cell lines. Prostate, 2007. 67(9): p. 955–67.

50. Ahmed, M., et al., Structural conversion of neurotoxic amyloid-beta(1-42) oligomers to fibrils. Nat Struct Mol Biol, 2010. 17(5): p. 561–7.

51. Sinha, S., D.H. Lopes, and G. Bitan, A key role for lysine residues in amyloid beta-protein folding, assembly, and toxicity. ACS Chem Neurosci, 2012. 3(6): p. 473–81.

52. Shukla, S., et al., Effect of Detergent Micelle Environment on P-glycoprotein (ABCB1)-Ligand Interactions. J Biol Chem, 2017.

53. Li, M.J., M. Guttman, and W.M. Atkins, Conformational dynamics of P-glycoprotein in lipid nanodiscs and detergent micelles reveal complex motions on a wide time scale. J Biol Chem, 2018. 293(17): p. 6297–6307.

54. Loo, T.W., et al., The ATPase activity of the P-glycoprotein drug pump is highly activated when the N-terminal and central regions of the nucleotide-binding domains are linked closely together. J Biol Chem, 2012. 287(32): p. 26806–16.

55. Griciuc, A., et al., Alzheimer’s disease risk gene CD33 inhibits microglial uptake of amyloid beta. Neuron, 2013. 78(4): p. 631–43.

56. Bradshaw, E.M., et al., CD33 Alzheimer’s disease locus: altered monocyte function and amyloid biology. Nat Neurosci, 2013. 16(7): p. 848–50.

57. Zhao, L., CD33 in Alzheimer’s Disease - Biology, Pathogenesis, and Therapeutics: A Mini-Review. Gerontology, 2019. 65(4): p. 323–331.

58. Brenn, A., et al., St. John’s Wort reduces beta-amyloid accumulation in a double transgenic Alzheimer’s disease mouse model-role of P-glycoprotein. Brain Pathol, 2014. 24(1): p. 18–24.

59. Morris, G.M., et al., AutoDock4 and AutoDockTools4: Automated docking with selective receptor flexibility. J Comput Chem, 2009. 30(16): p. 2785–91.

60. Dawson, R.J. and K.P. Locher, Structure of a bacterial multidrug ABC transporter. Nature, 2006. 443(7108): p. 180–5.

61. Sauna, Z.E. and S.V. Ambudkar, About a switch: how P-glycoprotein (ABCB1) harnesses the energy of ATP binding and hydrolysis to do mechanical work. Mol Cancer Ther, 2007. 6(1): p. 13–23.

62. Dawson, R.J. and K.P. Locher, Structure of the multidrug ABC transporter Sav1866 from Staphylococcus aureus in complex with AMP-PNP. FEBS Lett, 2007. 581(5): p. 935–8.

63. Dawson, R.J.P. and K.P. Locher, Structure of the multidrug ABC transporter Sav1866 from Staphylococcus aureus in complex with AMP-PNP. FEBS Lett, 2007. 581(5): p. 935–8.

64. Ward, A., et al., Flexibility in the ABC transporter MsbA: Alternating access with a twist. Proc Natl Acad Sci U S A, 2007. 104(48): p. 19005–10.

65. Ryan, T.M., et al., Ammonium hydroxide treatment of Abeta produces an aggregate free solution suitable for biophysical and cell culture characterization. PeerJ, 2013. 1: p. e73.

66. Martin, C., et al., The molecular interaction of the high affinity reversal agent XR9576 with P-glycoprotein. Br J Pharmacol, 1999. 128(2): p. 403–11.

67. Schindelin, J., et al., Fiji: an open-source platform for biological-image analysis. Nat Methods, 2012. 9(7): p. 676–82.

68. Schindelin, J., et al., The ImageJ ecosystem: An open platform for biomedical image analysis. Mol Reprod Dev, 2015. 82(7-8): p. 518–29.

69. Schneider, C.A., W.S. Rasband, and K.W. Eliceiri, NIH Image to ImageJ: 25 years of image analysis. Nat Methods, 2012. 9(7): p. 671–5.

70. Linkert, M., et al., Metadata matters: access to image data in the real world. J Cell Biol, 2010. 189(5): p. 777–82.

71. Delannoy, S., et al., Nucleotide binding to the multidrug resistance P-glycoprotein as studied by ESR spectroscopy. Biochemistry, 2005. 44(42): p. 14010–9.

72. Hoffman, A.D., I.L. Urbatsch, and P.D. Vogel, Nucleotide binding to the human multidrug resistance protein 3, MRP3. Protein J, 2010. 29(5): p. 373–9.

73. Denisov, I.G. and S.G. Sligar, Nanodiscs in Membrane Biochemistry and Biophysics. Chem Rev, 2017. 117(6): p. 4669–4713.

