## Supplemental Information for "Transport of Alzheimer’s Associated Amyloid-β Catalyzed by P-glycoprotein"

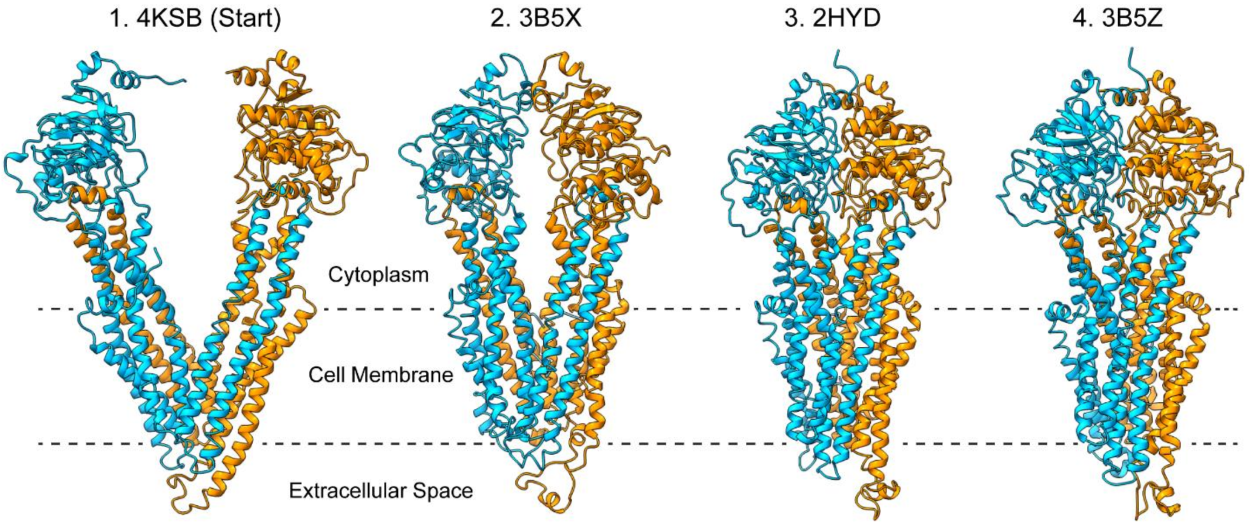


**Supplemental Figure 1. Conformational States of P-glycoprotein used in Targeted Molecular Dynamics Simulations.** Targeted Molecular Dynamics (TMD) simulations were performed as described in McCormick et al. 2015 using a dynamic model of human P-glycoprotein and several low energy conformations of P-gp homologues [35, 36]. Here we show the target names of the homologs and corresponding conformation of P-gp for each step in the putative catalytic cycle.

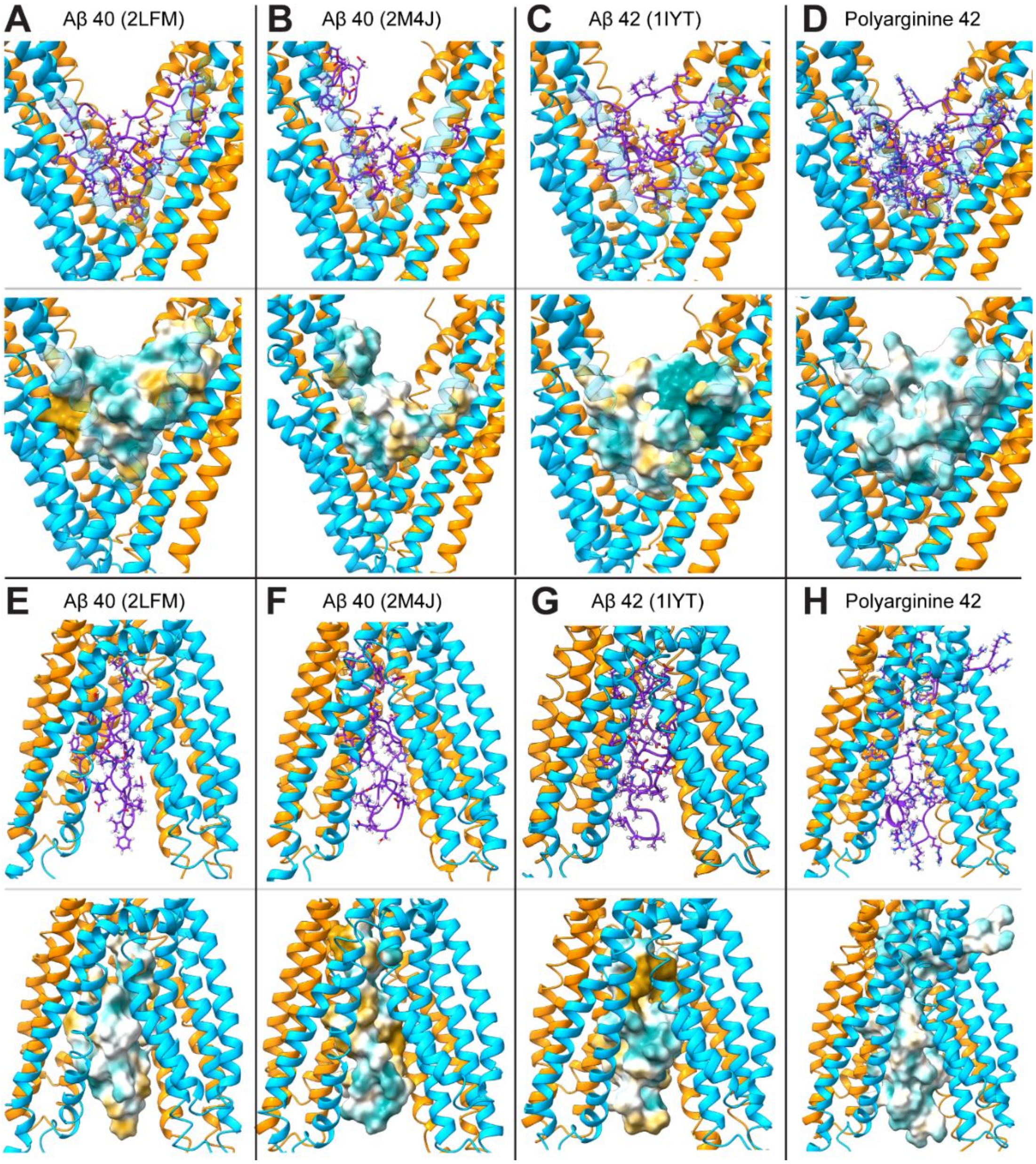


**Supplemental Figure 2. Aβ or Control Peptides at the First and Final Frames of Targeted Molecular Dynamics Experiments: A) Aβ40 (2LFM), B) Aβ40 (2M4J), C) Aβ42 (1IYT), D) Polyarginine 42**. Panels **A-D** are representative images of bound peptides at the start of molecular dynamics simulations. **(A)** 1IYT docked to the DBDs of P-gp with an estimated affinity of -7.2 kcal/mol. 1IYT is a solid state NMR structure of Aβ42 solved in an apolar microenvironment [4]. **B)** 2LFM is a partially folded Amyloid-β (Aβ) 40 structure that was solved in an aqueous environment [39]. 2LFM docked to the drug binding domains (DBDs) of P-glycoprotein (P-gp) with a predicted affinity of -7.2 kcal/mol. **C)** 2M4J is an Aβ40 fibril derived from brain tissue with Alzheimer's disease [38]. 2M4J docked to the DBDs of P-gp with an estimated affinity of -7.1 kcal/mol. Panels **E-H** are representative images at the end of a TMD simulation of the putative catalytic cycle. The bound peptide is shown in purple licorice representation or in surface representation colored for lipophilicity (teal = hydrophilic, gold = lipophilic, and white = neutral). The N- and C-terminal halves of P-gp are colored turquoise or orange.

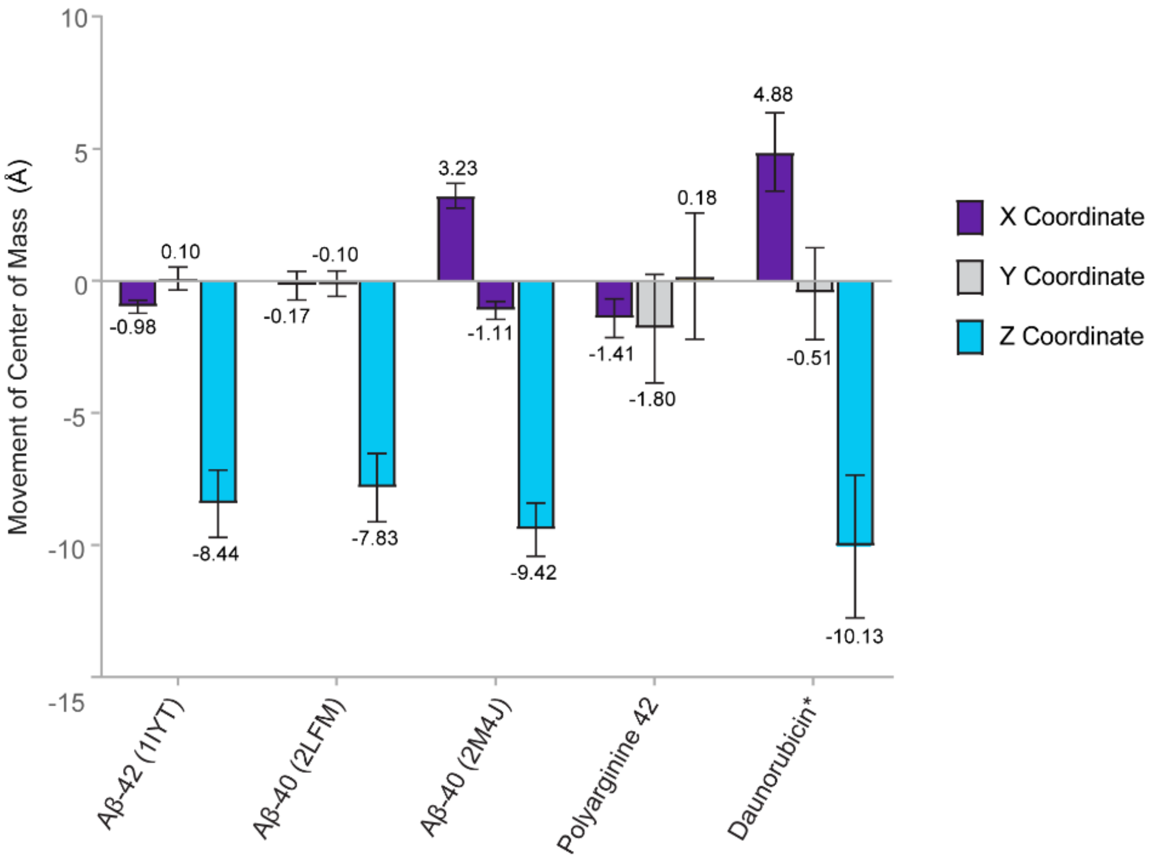


**Supplemental Figure 3. Total Movement of Amyloid-β, Polyarginine 42, or Daunorubicin through P-gp during a Putative Catalytic Cycle:** The center of mass of each ligand was calculated for each step of the simulations relative to the distance from the starting location in the X, Y, and Z axes. The plane of the membrane is parallel to the X and Y plane; movement through the membrane is oriented on the Z-axis. Distances are presented in Å. Six simulations were performed for each ligand; data represent the mean total transport distance ± one standard deviation from the mean.

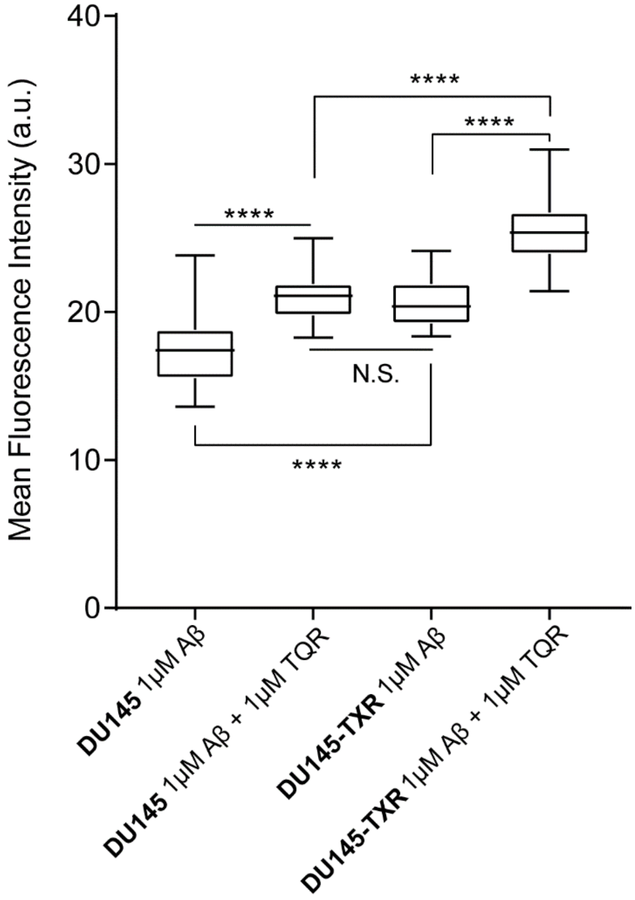


**Supplementary Figure 4. The Accumulation of fl-Aβ42 in DU145 and DU145-TXR Cancer Cells.** The intracellular fluorescence of paired chemotherapeutic sensitive/resistant cancer cell line (DU145 and DU145-TXR) was measured by confocal microscopy after incubation with 1 µM fluorescently labeled Aβ42 in the presence or absence of 1 µM Tariquidar (TQR). Statistical significance was determined using an unpaired T-test in Graphpad Prism; data are n = 24 images per treatment, two trials per treatment. Data are expressed as arbitrary units (a.u.) as calculated by the Integrated Density function of ImageJ [67-70].

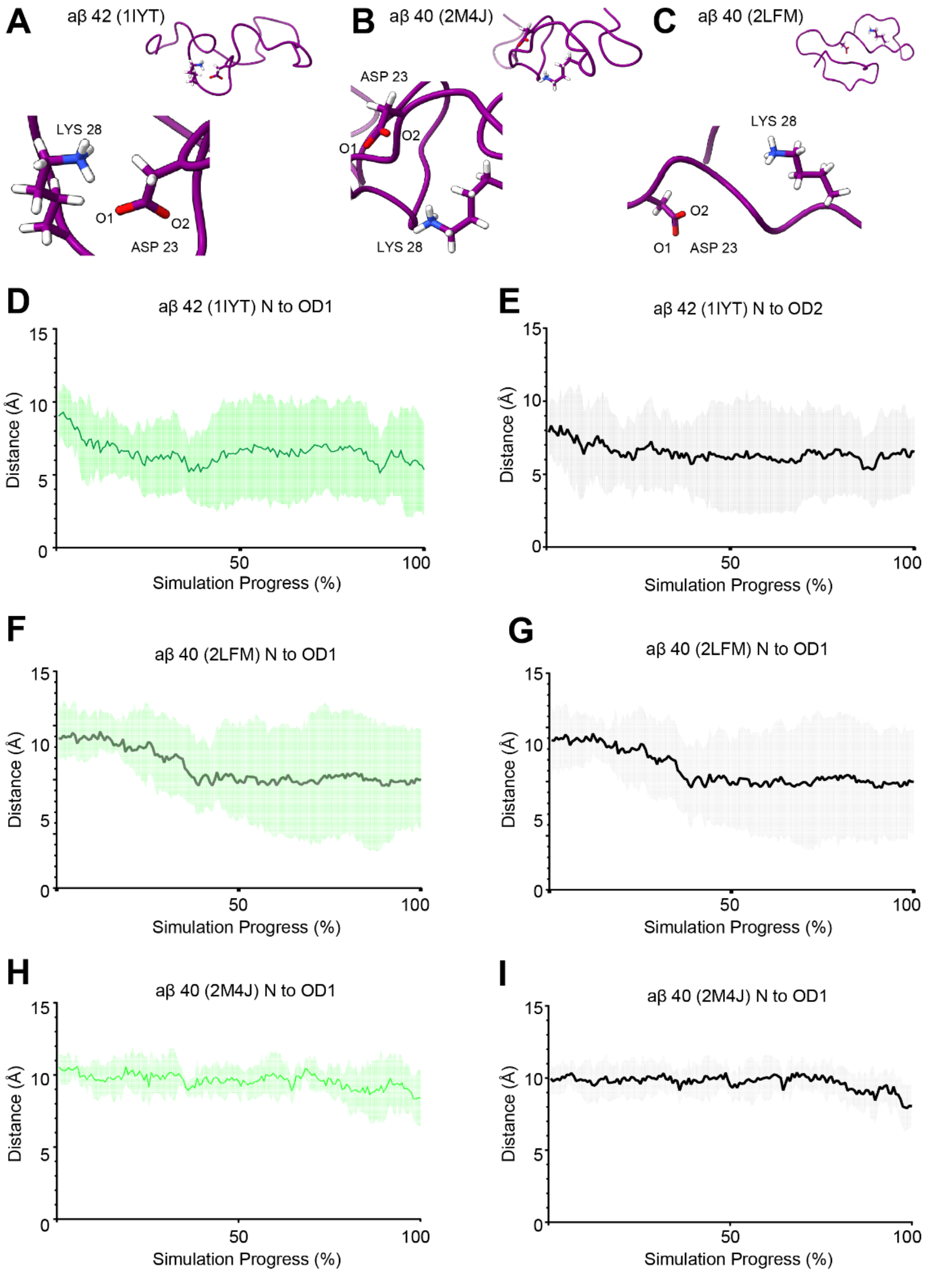


**Supplemental Figure 5. Distance between LYS28 and ASP23 of Aβ Monomers during Catalytic Transport Cycle of P-gp:** The Distance in Angstroms between the charged nitrogen (N) of LYS28 and either potentially charged oxygen (OD1, or OD2) of ASP23 in the respective Aβ peptides. Panels **A-C** show the position of K28 and D23 in each Aβ monomer. Graphs **D,E** show the mean distance in Angstroms between K28 and D23 in Aβ 42 (1IYT) during TMD simulations; graphs **F,G** show the mean distance in Angstroms between K28 and D23 in Aβ 40 (2LFM); graphs **H,I** show the mean distance in Angstroms between K28 and D23 in Aβ 40 (2M4J). Data represent the mean distance between the two selected atoms ± one standard deviation shown in colored shading (n = 6).

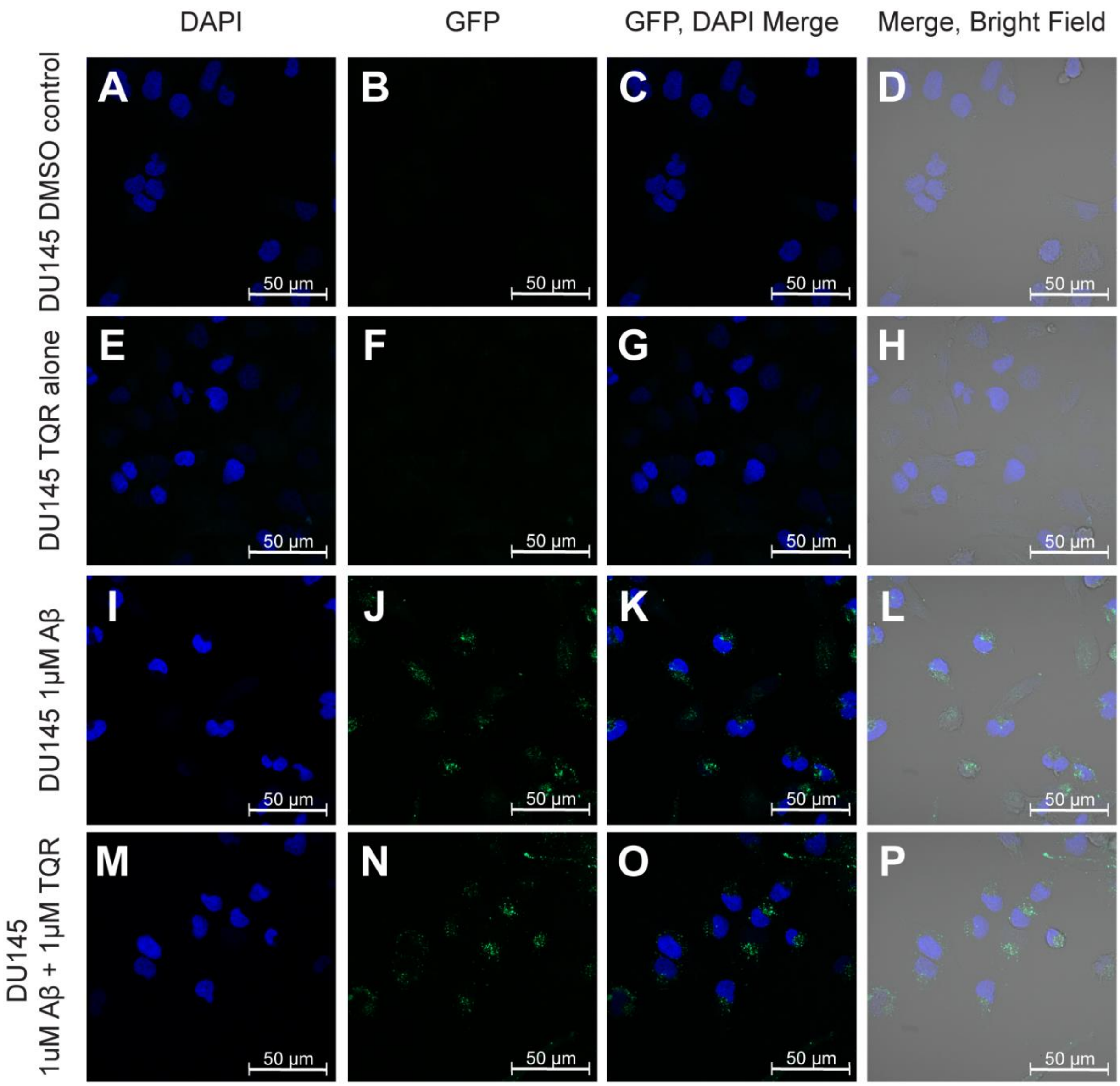


**Supplemental Figure 6. The Accumulation of fl-Aβ42 in DU145 Cancer Cells.** The intracellular fluorescence of chemotherapeutic sensitive DU145 cells was measured by confocal microscopy. Fluorescence was measured 24 hours after a 16 hour incubation with fluorescently labeled Aβ42 monomers. Representative Images **A-D** show DU145 treated with DMSO alone; representative images **E-H** show DU145 treated with 1 µM Tariquidar (TQR) alone; representative images **I**-L show DU145 treated with 1 µM Aβ42 alone; representative images **M-P** show DU145 treated with 1 µM Aβ42 and 1 µM TQR. Data are n = 24 images, two trials.

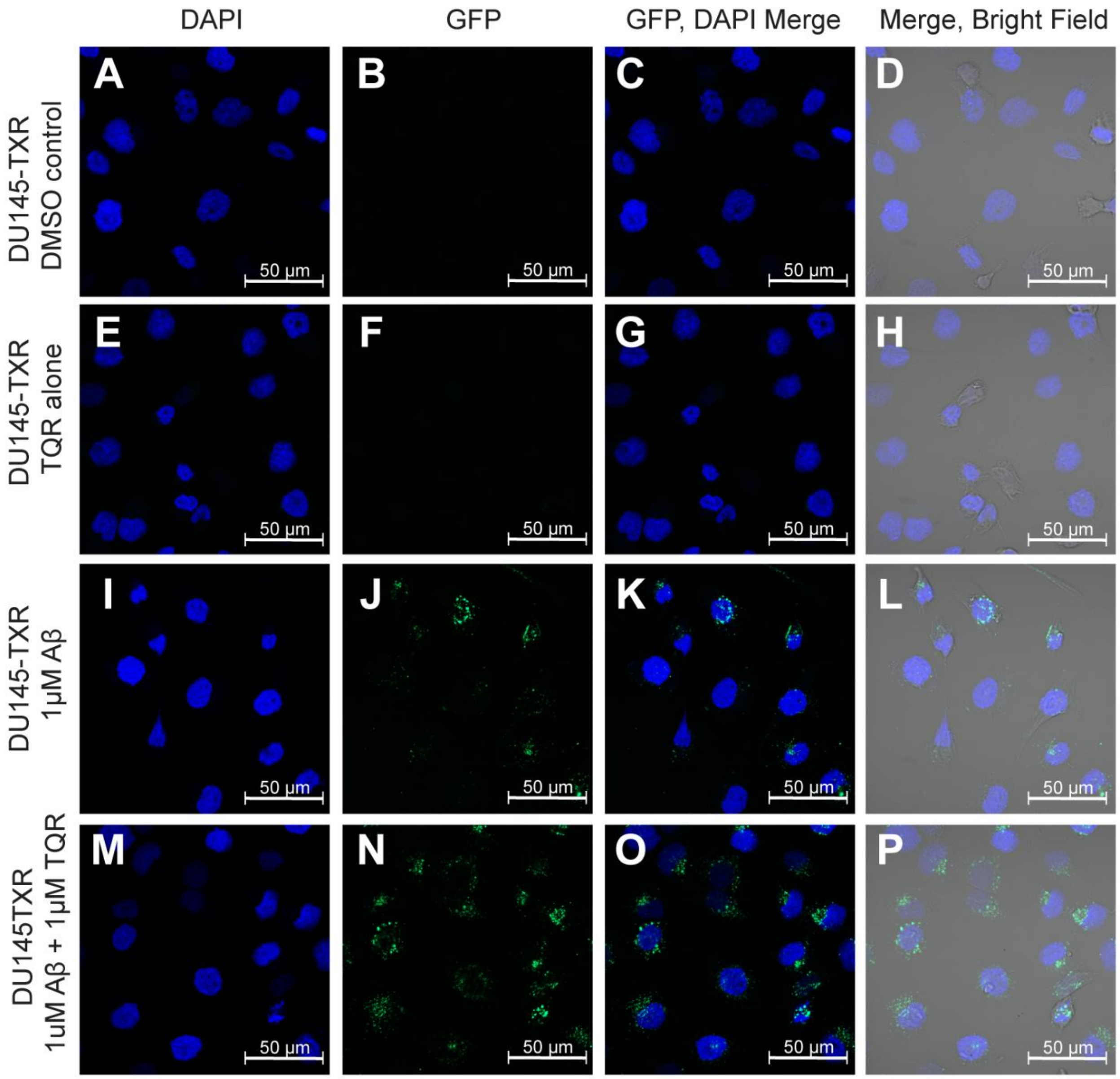


**Supplemental Figure 7. The Accumulation of fl-Aβ42 in DU145-TXR P-gp overexpressing Cancer Cells.** The intracellular fluorescence of chemotherapeutic resistant, P-gp overexpressing DU145-TXR cells was measured by confocal microscopy. Fluorescence was measured 24 hours after a 16 hour incubation with fluorescently labeled Aβ42 monomers. Representative Images A-D show DU145-TXR treated with DMSO alone; representative images E-H show DU145-TXR treated with 1 µM Tariquidar (TQR) alone; representative images I-L show DU145-TXR treated with 1 µM Aβ42 alone; representative images M-P show DU145-TXR treated with 1 µM Aβ42 and 1 µM TQR. Data are n = 24 images, two trials.

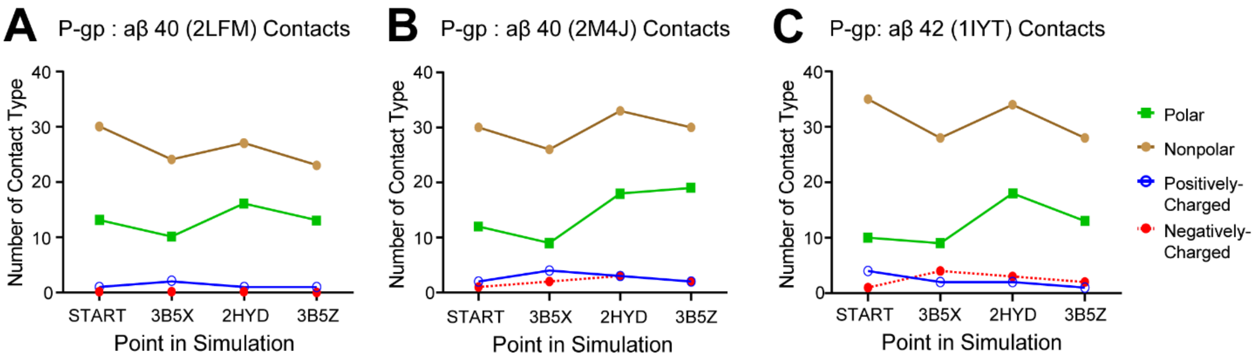


**Supplemental Figure 8. Number and Characteristics of Residue Contacts made by Aβ42 peptides to the Drug Binding Domains of P-gp during TMD Simulations.** Data are reported as the number and type of residues in the P-gp Drug Binding Domain with an α-carbon within 3 Å of the respective Aβ peptide. Residues are classified as Polar (SER, THR, CYS, ASN, GLN, TYR), Non-Polar (GLY, ALA, VAL, LEU, MET, ILE, PHE, PRO, TRP), Positively Charged (LYS, ARG, HIS), or Negatively Charged (GLU, ASP). Residues were included in the total count if the respective Aβ peptide interacted with the residue in 4/6 TMD simulations.

**Supplementary Table 1. Number and Type of Residue Contacts made by Aβ monomers with the Drug Binding Domains of P-gp during TMD simulations.** Data are reported as the number of residues in the P-gp Drug Binding Domain with an α-carbon within 3 Å of the Aβ peptide. Residues are classified as Polar (SER, THR, CYS, ASN, GLN, TYR), Non-Polar (GLY, ALA, VAL, LEU, MET, ILE, PHE, PRO, TRP), positively Charged (LYS, ARG, HIS), negatively Charged (GLU, ASP).

| | Residue Category | | | |
|---|---|---|---|---|
| aβ40 (2LFM) | Polar | Non-Polar | Positively Charged | Negatively Charged |
| 4KSB (Start) | 13 | 30 | 1 | 0 |
| 3B5X | 10 | 24 | 2 | 0 |
| 2HYD | 16 | 27 | 1 | 0 |
| 3B5Z (End) | 13 | 23 | 1 | 0 |
| aβ40 (2M4J) | Polar | Non-Polar | Positively Charged | Negatively Charged |
| 4KSB (Start) | 12 | 30 | 2 | 1 |
| 3B5X | 9 | 26 | 4 | 2 |
| 2HYD | 18 | 33 | 3 | 3 |
| 3B5Z (End) | 19 | 30 | 2 | 2 |
| aβ42 (1IYT) | Polar | Non-Polar | Positively Charged | Negatively Charged |
| 4KSB (Start) | 10 | 35 | 4 | 1 |
| 3B5X | 9 | 28 | 2 | 4 |
| 2HYD | 18 | 34 | 2 | 3 |
| 3B5Z (End) | 13 | 28 | 1 | 2 |

**Supplementary Table 2. Transport of Aβ and Daunorubicin by P-gp According to Structural Transition.** Distances are reported in Angstroms relative to the center of mass coordinates at the start of the simulation, and are reported as mean ± standard deviation, n = 6.

| | Start (4KSB) | Δ 4KSB -> 3B5X | 3B5X | Δ 3B5X -> 2HYD | 2HYD | Δ 2HYD- > 3B5Z | End (3B5Z) |
|---|---|---|---|---|---|---|---|
| Aβ 42 | 0.00 | -1.55 ± .054 | -1.55 ± .054 | -4.44 ± 1.68 | -5.98 ± 1.80 | -2.45 ± 1.06 | -8.44 ± 1.27 |
| Aβ 40 2M4J | 0.00 | -3.11 ± 0.67 | -3.11 ± 0.67 | -3.77 ± 0.96 | -6.88 ± 1.16 | -2.54 ± 0.74 | -9.42 ± 1.01 |
| Aβ 40 2LFM | 0.00 | -1.04 ± 0.98 | -1.04 ± 0.98 | -2.72 ± 1.88 | -3.76 ± 1.84 | -4.06 ± 2.76 | -7.83 ± 1.29 |
| POLY42 | 0.00 | 5.11 ± 1.31 | 5.11 ± 1.31 | -3.01 ± 1.14 | 2.11 ± 2.22 | -1.89 ± 0.99 | 0.21 ± 2.58 |
| DAU | 0.00 | -2.85 ± 2.21 | -2.85 ± 2.21 | -2.70 ± 2.74 | -5.54 ± 3.41 | -4.59 ± 2.24 | -10.13 ± 2.72 |

**Supplementary Table 3.** The mean fluorescence intensity of paired chemotherapeutic sensitive/resistant cancer cell line (DU145 and DU145-TXR) as measured by confocal microscopy after incubation with 1 µM fluorescently labeled Aβ42 in the presence or absence of 1 µM Tariquidar (TQR). Statistical significance was determined using an unpaired T-test in Graphpad Prism; data are n = 24 images per treatment, two trials per treatment. Data are expressed as arbitrary units (a.u.) as calculated by the Integrated Density function of ImageJ [67-70]. The table shows the mean fluorescence intensity (a.u.) of per image ± one standard deviation from the mean.

| Cell Line | Treatment | Mean Fluorescence Intensity (a.u.) ± standard deviation |
|---|---|---|
| DU145 | 1µM Aβ | 17.65 ± 2.42 |
| DU145 | 1µM Aβ + 1µM TQR | 20.96 ± 1.62 |
| DU145-TXR | 1µM Aβ | 20.60 ± 1.62 |
| DU145-TXR | 1µM Aβ + 1µM TQR | 25.57 ± 2.38 |
